# Ancient dental calculus preserves signatures of biofilm succession and inter-individual variation independent of dental pathology

**DOI:** 10.1101/2022.04.25.489366

**Authors:** Irina M. Velsko, Lena Semerau, Sarah A. Inskip, Maite Iris García-Collado, Kirsten Ziesemer, Maria Serrano Ruber, Luis Benítez de Lugo Enrich, Jesús Manuel Molero García, David Gallego Valle, Ana Cristina Peña Ruiz, Domingo C. Salazar García, Menno L.P. Hoogland, Christina Warinner

## Abstract

Dental calculus preserves oral microbes, enabling comparative studies of the oral microbiome and health through time. However, small sample sizes and limited dental health metadata have hindered health-focused investigations to date. Here we investigate the relationship between tobacco pipe smoking and dental calculus microbiomes. Dental calculus from 75 individuals from the 19th century Middenbeemster skeletal collection (Netherlands) were analyzed by metagenomics. Demographic and dental health parameters were systematically recorded, including the presence/number of pipe notches. Comparative data sets from European populations before and after the introduction of tobacco were also analyzed. Calculus species profiles were compared with oral pathology to examine associations between microbiome community, smoking behavior, and oral health status. The Middenbeemster individuals exhibited relatively poor oral health, with a high prevalence of periodontal disease, caries, heavy calculus deposits, and antemortem tooth loss. No associations between pipe notches and dental pathologies, or microbial species composition, were found. Calculus samples before and after the introduction of tobacco showed highly similar species profiles. Observed inter-individual microbiome differences were consistent with previously described variation in human populations from the Upper Paleolithic to the present. Dental calculus may not preserve microbial indicators of health and disease status as distinctly as dental plaque.

**Research Highlights:**

- No associations between calculus species profiles and oral health metrics were detected in a single large population
- A minority of individuals have a dental calculus species profile characterized by low levels of *Streptococcus* and high levels of anaerobic taxa

## Introduction

Dental calculus is a mineralized form of dental plaque that forms on the surface of teeth during life and persists in the archaeological record. Diverse microremains and biomolecules, including DNA, protein, and metabolites, are preserved within ancient dental calculus (Adler et al., 2013; Hardy et al., 2016; Hendy et al., 2018; Salazar-García et al., 2021; Velsko et al., 2019, 2017; Warinner et al., 2014) and can be used to study oral microbial ecology and evolution through time (Fellows Yates et al., 2021b; Granehäll et al., 2021; Ottoni et al., 2021), as well as provide evidence of past human activities (Radini et al., 2019). The majority of biomolecules present in calculus derive from dental plaque bacteria, and there is great interest in determining the feasibility of using these microbes to indirectly trace evidence of human behavioral or lifestyle changes and their impact on health through deep time. Certain past activities, such as the rise and spread of tobacco smoking during the European Colonial period, can be difficult to detect directly but likely had important health consequences. Examining dental plaque communities via dental calculus may enable the detection of tobacco use and its growing impact on oral health, but to date this topic has not been extensively explored.

Tobacco was introduced to Europe from the Americas at the turn of the 15th century, initially as a medical remedy (Goodman, 1993). However, by the late 16th century, tobacco smoking had become a popular social and leisure activity, particularly in Western Europe (Brongers, 1964; Gately, 2001; Goodman, 1993). Archaeologically, it has been possible to detect pipe smoking through the identification of so-called dental ‘pipe notches’. Pipe notches are areas of dental abrasion caused by habitually clenching a clay pipe between the anterior teeth (Figure 1B, C, Supplemental Figure S1). Multiple studies of these features in 17th-19th century European populations have shown that pipe smoking was a predominantly male activity which varied in popularity over time but increasingly became linked to lower socioeconomic status in the 18th and 19th century (Geber and Murphy, 2018; Inskip et al., n.d.; Kvaal and Derry, 1996; Veselka, 2016; Walker and Henderson, 2010). These trends correspond to historical records about tobacco use and smoking in England and the Netherlands (Goodman, 1993; Hughes, 2003; Tullett, 2019).

**Figure 1.**
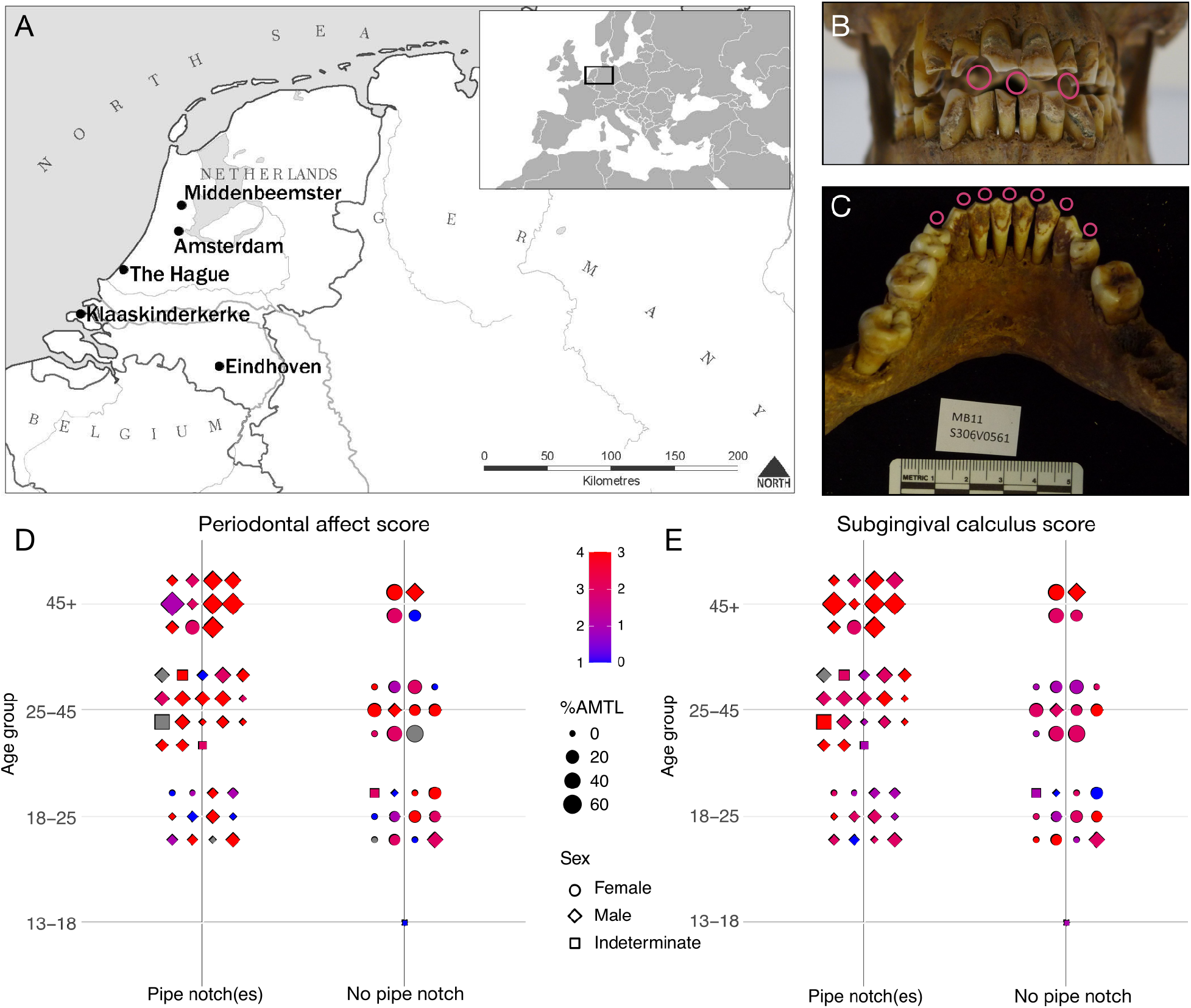
The Middenbeemster collection. **A.** Map locating the region of Middenbeemster in the Netherlands and Europe. **B.** Male individual from Middenbeemster with three prominent pipe notches in the anterior dentition indicated by hollow pink dots. **C.** Mandible of a male individual S306V0561 with at least 7 pipe notches on the anterior dentition, indicated by hollow pink dots. **D,E.** Dental pathology of Middenbeemster individuals in relation to age, sex, antemortem tooth loss (AMTL) and presence of pipe notches, points colored by **D.** periodontal affect score (1-4), or **E.** subgingival calculus score (0-3). Gray indicates no data. Photo credit: Sarah A. Inskip.

Studies of present-day tobacco users have reported a negative relationship between tobacco use and oral health (Albandar et al., 2000; Axelsson et al., 1998; Huang and Shi, 2019). Tobacco users have been shown to have more severe dental and periodontal pathologies, including caries and tooth loss (Heng et al., 2006; Vellappally et al., 2007), periodontal disease (Bergstrom, 2014; Pejčić et al., 2007; Sreedevi et al., 2012), and calculus accumulation (Albandar et al., 2000; Baljoon et al., 2005; Martinez-Canut et al., 1995; Sreedevi et al., 2012). Chemicals inhaled in tobacco smoke appear to affect oral microbes and promote a more pathogenic community, such that multiple studies have found that smokers have distinct dental plaque microbial communities compared to non-smokers (Al Bataineh et al., 2020; Kumar et al., 2011; Mason et al., 2015; Moon et al., 2015; Yang et al., 2019). However, the extent to which these differences are also present and persist in dental calculus, which represents a more mature biofilm than dental plaque (Velsko et al., 2019), is not known. Potentially, dental calculus may preserve these distinctions, and if so, would offer the possibility of identifying heavy smokers in the past, including in cases where abrasive clay pipes were not used.

Here we investigated the dental calculus microbial communities associated with pipe smoking in historic European populations, particularly focusing on 19th century individuals from Middenbeemster (MID), the Netherlands (Figure 1A). Contrary to expectations, we found that there are minimal differences in the dental calculus microbial structure of individuals with evidence of heavy pipe smoking (pipe notches) and those without, despite evidence for differences in skeletal markers of oral disease between those groups. Moreover, we find that through comparative analysis with additional Medieval, Industrial, and Modern (present-day) individuals, this pattern holds more broadly, with no discernible differences in overall microbial community profiles between individuals who lived before and after the introduction of tobacco to Europe.

A minority of MID individuals (9/72, 12%) exhibited a distinct oral microbial profile characterized by a greatly reduced number of early colonizer taxa, including *Streptococcus*. A similar oral microbial profile at a similar prevalence (11%) was also observed among a diverse range of humans from Europe and Africa from the Paleolithic onwards (Fellows Yates et al., 2021b). Through analysis of the MID assemblage and a set of densely sampled Chalcolithic dentitions (Fagernäs et al., 2022), we demonstrate that this pattern is not associated with dental pathology, but may instead reflect long-term standing variation in patterns of plaque biofilm development among humans.

## Results

### Dentition

#### Demography and pipe use

Dental calculus was collected from a total of 75 individuals in the Middenbeemster collection, including 40 males, 25 females, and 5 individuals of unknown sex. A total of 66 of individuals could be aged (Table 1). Of the 25 females, 24 could be attributed an age, while for the 40 males, 38 could be provided an age estimate. In general there are proportionally more females in the young category, and more males in the old category, which needs to be considered when assessing oral pathology.

**Table 1.** Age and sex distribution of individuals in the study sample.

| Age (yearss) | Number Males<sup>1</sup> | Number Females<sup>1</sup> | Number Unknown sex<sup>1</sup> |
| --- | --- | --- | --- |
| Young (18-25) | 15 (22.7%) | 13 (19.7%) | 1 (1.5%) |
| Middle (25-45) | 14 (21.2%) | 8 (12.1%) | 3 (4.5%) |
| Old (>45) | 9 (13.6%) | 3 (4.5%) | 0 |
| Unknown | 2 | 1 | 1 |
| **Total** | **40** | **25** | **5** |

<sup>1</sup>Percent indicates percent of total aged individuals (66).

The dentition was sufficiently intact to score the presence or absence of pipe notches in 70 individuals (Table S2, Figure 1B). We observed a strong relationship between biological sex and pipe smoking, with most individuals bearing pipe notches being male. Only 3 of 25 females had pipe notches (12%). Two of these women had one notch, and the other had two. Conversely, most men exhibited pipe notches (88%; 35/40). Of the five men who did not have a notch, two of these are young males and three are middle-age males. Of those with notches, 75% had at least two notches. The maximum number of notches observed among men was 7 (Figure 1C). The pattern of pipe notches in males suggests that habitual pipe smoking started from a young age. Furthermore, based on this data, pipe smokers were slightly older on average than non-smokers.

### Oral pathologies by pipe use status

While there was no difference in the incidence of caries between individuals with and without pipe notches, individuals without pipe notches had a higher percentage of teeth with caries (26%) compared to pipe-users (15.1%) (t=2.1194, df=66, p = 0.038), and more non-users had gross caries (50%) than pipe-users (27.5%). However, there was a trend for more of the pipe users to have lost teeth antemortem (80.5%) than non-users (67.9%), possibly confounding our observations (Table S3).

Early stages of periodontal disease (stage 2) was observable in almost all individuals. As such, we assessed whether there were differences in more severe stages of periodontal disease between pipe-users and non-users (stages 3 and 4). More pipe-users had periodontal disease at stage 3 or 4 (87.8%) than non-users (64.3%), a difference that was statistically significant by chi-squared test (p = 0.035, n = 69). In terms of periapical lesions, more pipe users were affected (45%) compared to non-users (35.7%), and had more positions affected (3.4) compared to non-users (2.4). Additionally, dental calculus was almost ubiquitous in the sample collection, so we assessed how many teeth were affected and the general level of severity in the mouth. Pipe users had a greater proportion of teeth with dental calculus (73.5%) compared to non-users (58.6%), and had a greater build-up of both supra-and subgingival calculus.

To better understand patterns of oral pathologies, we assessed the relationship between each condition and age. There was a positive relationship between prevalence and age for all oral pathologies with the exception of caries and gross caries. Caries prevalence was similar in all age groups, while gross caries became less common with age. While this may seem counterintuitive, caries is a leading cause of tooth loss, and AMTL prevalence was highest in the oldest age category. The lower prevalence of caries, especially of gross caries, in the oldest category is likely due to the high degree of observed tooth loss.

Figures 1D and E graphically depict the severity of three oral pathologies that have been linked to smoking in present-day modern populations, including periodontal disease, antemortem tooth loss (AMTL) and subgingival calculus (Al-Zahrani et al., 2021; Bergström, 2005; Eke et al., 2015; Kowalski, 1971; Lee et al., 2022), by age and sex. For all three oral pathologies, middle-age and old-age adults tended to have more severe manifestations of the conditions, with nearly all individuals in the 45+ years category having periodontal disease scores of 3 or 4, calculus scores of 2 or 3, and high rates of tooth loss (mean 28% of teeth per individual, compared to mean 8.8% for all other age groups). As such, it is evident that there is a strong positive relationship between the severity of oral pathologies and age but not sex (Figure 1D,E).

### Calculus preservation assessment

To explore whether microbial signatures of smoking could be detected from 18th century Europe, we analzyed dental calculus samples from Middenbeemster (MID), the Netherlands, and Convento de los Mercedarios de Burtzeña (CMB), Spain. Dental calculus microbial community preservation was high at both sites (Supplemental Figures S2, S3), with all but 3 MID samples passing preservation thresholds. Eight MID samples did not have sufficient metadata for further comparisons and were excluded from all microbiome analysis. After filtering for preservation and metadata completeness, 73 samples were used in downstream analyses (MID = 65, CMB = 8).

### Species profile differences

We first wanted to determine if there are broadly discernable differences in calculus species profiles related to smoking evidence within the MID and CMB populations. To compare microbial profile differences related to smoking status, individuals were classified as heavy smokers if their dentition showed one or more pipe notches, or light/non-smokers if there was no sign of pipe notches in their dentition. Beta-diversity analysis was performed with a principal components analysis (PCA) to compare species profiles. Canonical correlation analysis was used to assess correlations between sample metadata, including oral pathology, laboratory outcomes, and sequencing analysis, as well as between sample metadata and principal component loadings (Figure 2A).

**Figure 2.**
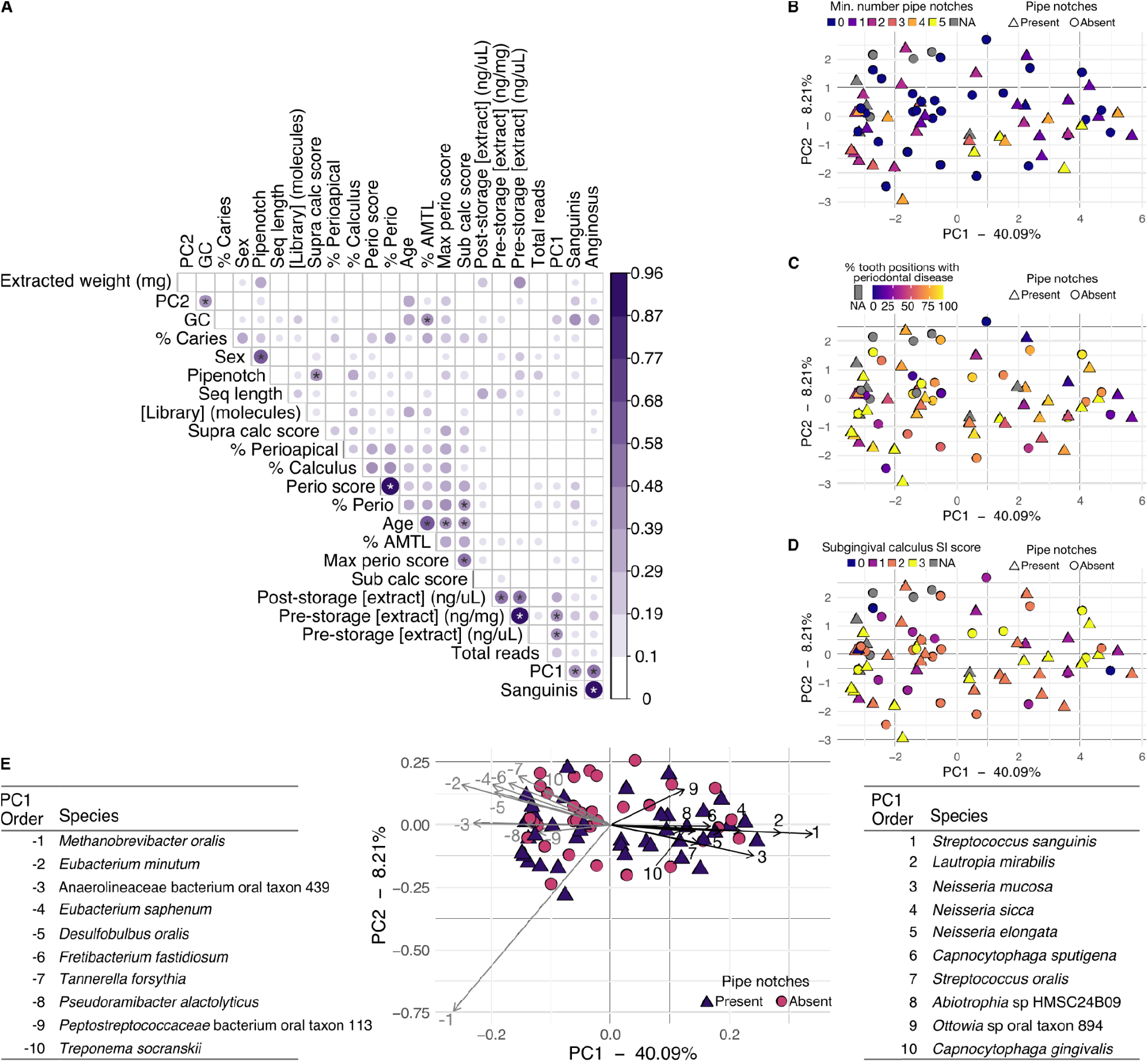
Microbial community diversity and correlations with oral pathology and laboratory work metadata. **A.** Canonical correlations (CC) for Middenbeemster samples between metadata categories and principal components loadings. Significance tests were performed with a Pearson correlation test. The size and color of the dots corresponds to the CC value, which does not determine the direction of the correlation (positive or negative. Hence all CC values are positive). Correlations ≥ 0.4 have significance indicated with stars. * p ≤ 0.001. B-E PCA based on species composition of MID and CMB samples colored by **B.** Minimum number of pipe notches. **C.** Percent of tooth positions with periodontal disease. **D.** Subgingival calculus SI score. Samples from CMB are colored gray in B-D because due to the fragmented nature of the skeletons, the same metadata could not be collected (see Supplemental Figure S1). **E.** PCA based on species composition of MID and CMB samples colored by pipe notch presence, including a bi-plot indicating the loadings of 10 species with strongest positive and negative PC1 loadings, with the species indicated by number corresponding to the strength of loading (1/-1 is strongest, 10/-10 is weakest). The species are listed in tables to the left and right of the plot, ordered by decreasing strength of the loading. Metadata shown in (**A**): **extracted weight (mg)** - weight of calculus used in extraction; **PC2** - PC2 loadings; **GC** - library average GC content; **% Caries** - % of teeth with caries; **Sex** - estimated biological sex; **Pipenotch** - pipe notch present; **Seq length** - library average sequence length; **[Library] (molecules)** - total DNA molecules in the library (x 106); **Supra calc score** - subragingival calculus SI score; **% Perioapical** - % of teeth with perioapical lesions; **% Calculus** - % of teeth with calculus; **Perio score** - average periodontitis score; **% Perio** - % of teeth with periodontal disease; **% AMTL** - % of teeth lost ante-mortem; **Max perio score** - maximum periodontitis score; **Sub calc score** - subgingival calculus SI score; **Post-storage [extract]** (ng/uL) - extract DNA concentration after storage; **Pre-storage [extract] (ng/mg)** - extract DNA concentration directly after extraction; **Pre-storage [extract] (ng/uL)** - extract DNA concentration directly after extraction; **Total reads** - total reads in the library after quality-trimming and merging; **PC1** - PC1 loadings; **Sanguinis** - proportion of total reads that were assigned to a species in the Sanguinis *Streptococcus* group; **Anginosus** - proportion of total reads that were assigned to a species in the Anginosus *Streptococcus* group.

Strong correlations were found between sex and the presence of pipe notches, and between individual age at death and the following oral pathologies: percent antemortem tooth loss, percent of teeth with periodontal disease, maximum periodontal disease score, and subgingival calculus score. PC1 loadings were found to be correlated with extracted DNA concentration, and the proportions of *Streptococcus* taxa belonging to the Sanguinis and Anginosus groups in the samples, while PC2 loadings were correlated with average library GC content. However, no clustering with respect to microbial community composition and evidence of smoking status in PCA was observed, such as minimum number of pipe notches (Figure 2B), or periodontal health, such as percent of tooth positions with periodontal disease (Figure 2C), subgingival calculus score (Figure 2D), and others (Supplemental Figure S5). This suggests that these health metrics are not shaping the species profiles in this dataset, and other factors may be involved.

To understand which species were driving the sample plotting patterns, we examined the top ten species with the strongest positive and negative loadings in PC1 (Figure 2E). We found the species separating samples along PC1 have different environmental niches. The top ten species with strongest negative loadings are largely anaerobic taxa that are dominant in mature oral biofilms, including those in the genera *Methanobrevibacter*, *Eubacterium*, *Desulfobulbus*, *Fretibacterium*, and *Tannerella*. In contrast, the top ten species with strongest positive loadings are largely aerobic or facultative taxa that grow well in the presence of oxygen and are dominant in early dental biofilm formation, including those in the genera *Streptococcus*, *Neisseria*, and *Capnocytophaga*. The samples with high positive PC1 loadings, indicating a strong presence of oxygen-tolerant, early colonizer taxa, also show a higher proportion of ‘plaque’ in the SourceTracker plots (Supplemental Figure S3), supporting that they have a species profile that appears to have calcified at an earlier stage of development.

### Microbial functional profile

Although we found no differences in the species profiles between heavy- and light/non-smokers, and few associations between the species profiles and any metadata we collected from MID and CMB, we next investigated microbial gene content in the calculus from these populations. The extent to which inferred metabolic activity from ancient metagenomic data may reflect biofilm activity is an open area of investigation. Clinical periodontal microbiome research has reported distinctive gene expression profiles between dental plaque samples on teeth with and without periodontitis, even when there were not distinctive taxonomic differences (Duran-Pinedo et al., 2014; Yost et al., 2017, 2015). We used HUMAnN3 (Beghini et al., 2021) to infer the metabolic pathways present based on gene content in the MID and CMB samples, and performed PCA and canonical correlation analysis to assess the associations between sample metadata and the inferred metabolic potential of these calculus microbial communities (Figure 3).

**Figure 3.**
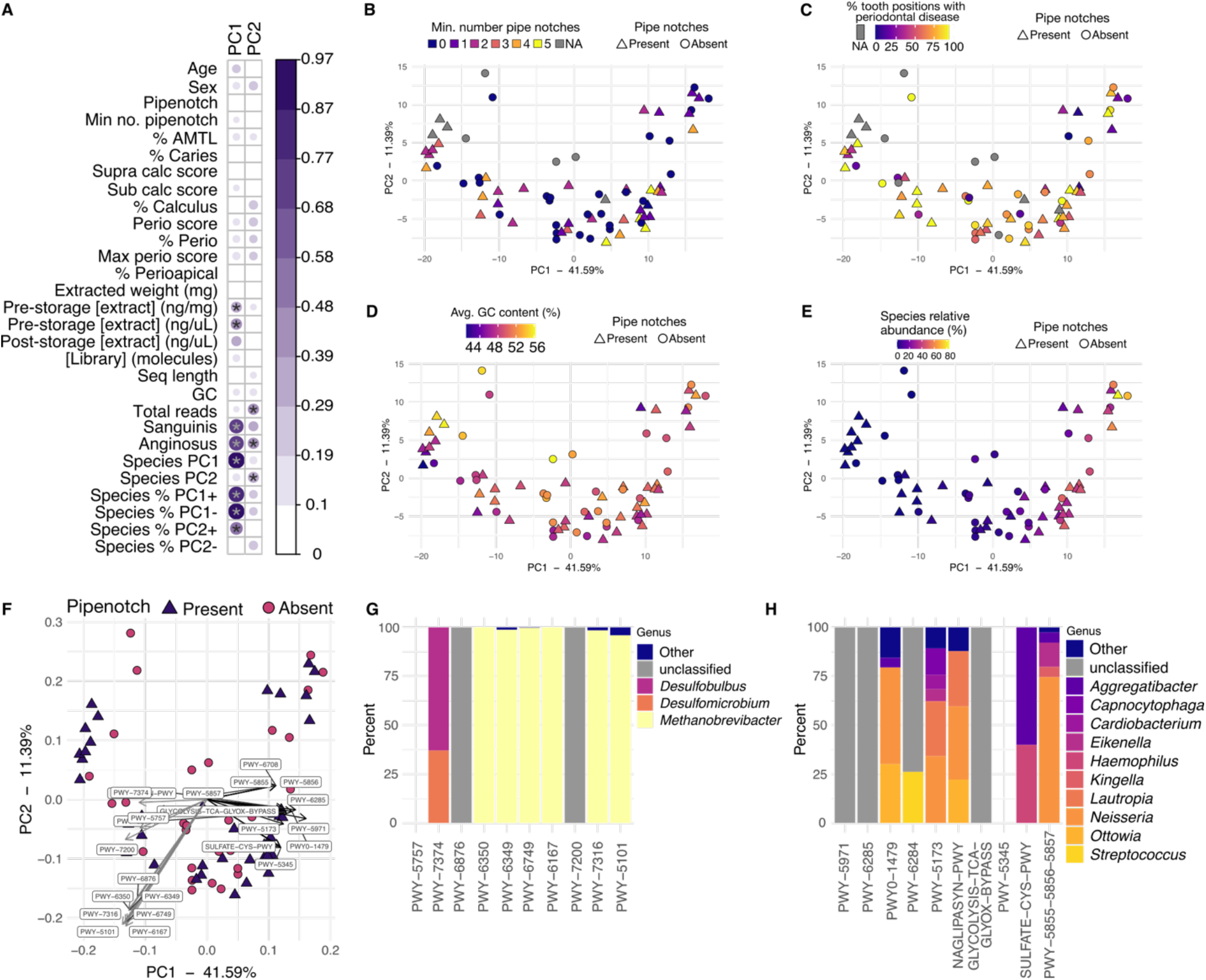
Metabolic pathway analysis and correlations. **A.** Canonical correlations (CC) for MID and CMB samples between metadata categories and principal component loadings for a PCA based on pathway abundance. Significance tests were performed with a Pearson correlation test. The size and color of the dots corresponds to the CC value, which does not determine the direction of the correlation (positive or negative. Hence all CC values are positive). Correlations ≥ 0.4 have significance indicated with stars. * p ≤ 0.001. **B-E** PCA plot based on pathway abundance in MID and CMB samples colored by **B.** Minimum number of pipe notches, **C.** Percent of teeth with periodontal disease, **D.** Average GC content (%), and **E.** Relative abundance of the 10 species with strongest PC1 positive loadings from the species PCA in Figure 2E. In (**C**) Samples from CMB are colored gray because due to the fragmented nature of the skeletons, the same metadata could not be collected (see Supplemental Figure S1). **F.** PCA biplot showing the 20 pathways with strongest loadings in PC1 (10 highest positive loadings, 10 highest negative loadings). **G** and **H** show the percent of each pathway contributed by species in the indicated genera. **G.** Ten pathways with strongest positive PC1 loadings. **H.** Ten pathways with strongest negative PC1 loadings. PWY-5855, PWY-5856, and PWY-5857 all have the same PC1 loading, and are contributed by the same proportions of the same species. All genera for which the total contribution was < 5% are grouped together as Other. The empty places for PWY-5757 in D and PWY-5345 in G indicate that HUMAnN3 was not able to attribute these pathways to specific species. Metadata shown in **A**: **PC1** - PC1 loadings; **PC2** - PC2 loadings; **Age** - estimated age at death; **Sex** - estimated biological sex; **Pipenotch** - one or more pipe notches present; **Min no. pipenotch** - minimum number of pipe notches; **% AMTL** - % of teeth lost ante-mortem; **% Caries** - % of teeth with caries; **Supra calc score** - subragingival calculus SI score; **Sub calc score** - subgingival calculus SI score; **% Calculus** - % of teeth with calculus; **Perio score** - average periodontitis score; **% Perio** - % of teeth with periodontal disease; **Max perio score** - maximum periodontitis score; **% Perioapical** - % of teeth with perioapical lesions; **Extracted weight (mg)** - weight of calculus used in extraction; **Pre-storage [extract] (ng/mg)** - extract DNA concentration directly after extraction; **Pre-storage [extract] (ng/uL)** - extract DNA concentration directly after extraction; **Post-storage [extract] (ng/uL)** - extract DNA concentration after storage; **[Library] (molecules)** - total DNA molecules in the library (x 10^6^); **Seq length** - library average sequence length; **GC** - library average GC content; **Total reads** - total reads in the library after quality-trimming and merging; **Sanguinis** - proportion of total reads that were assigned to a species in the Sanguinis *Streptococcus* group; **Anginosus** - proportion of total reads that were assigned to a species in the Anginosus *Streptococcus* group; **Species PC1** - Sample loading in PC1 from the PCA based on the MetaPhlAn3 species table; **Species PC2** - Sample loading PC2 from the PCA based on the MetaPhlAn3 species table; **Species % PC1+** - percent of 10 species with strongest PC1+ loadings in the species-based PCA out of total species; **Species % PC1-** - percent of 10 species with strongest PC1-loadings in the species-based PCA out of total species; **Species % PC2+** - percent of 10 species with strongest PC2+ loadings in the species-based PCA out of total species; **Species % PC2-** - percent of 10 species with strongest PC1-loadings in the species-based PCA out of total species.

Similar to the species-based canonical correlations, we found few strong correlations (>0.4) between the PCA principal component loadings and our sample metadata (Figure 3A), with the strongest being between the proportion of Sanguinis and Anginosus group streptococci in the samples. The principal component loadings of PC1 were also strongly correlated with the PC1 loadings from the species-based PCA, indicating that the sample loadings are shaped by similar factors in both the taxonomic and metabolic pathway PCA. Plots of PCAs revealed no distinctive clustering of the samples based on the minimum number of pipe notches (Figure 3B), the percent of teeth with periodontal disease (Figure 3C), or the library average GC content (Figure 3D). However, the samples with the highest positive PC1 loadings had the highest percentage of species with strongest positive PC1 loadings in the species PCA (Figure 3E, Figure 2E). Finally, we investigated the species that contribute to the 10 metabolic pathways with the strongest positive and negative loadings in PC1 (Figure 3F). The pathways with strongest negative PC1 loadings are mainly contributed by late colonizer, anaerobic taxa in the genera *Methanobrevibacter*, *Desulfobulbus*, and *Desulfomicrobium* (Figure 3G), while the pathways with the strongest positive PC1 loadings are contributed by a variety of aerobic and facultative species in the genera including *Eikenella*, *Haemophilus*, *Kingella*, *Lautropia*, *Neisseria*, *Ottowia* (Figure 3H). The difference in species contributing to pathways separating samples along PC1 reflects the gradient of taxa in the species-based PCA (Figure 2E), where samples at one end are characterized by a strong presence of early-colonizer taxa, while those at the other end are characterized by a strong presence of late-colonizer taxa. Both the species-based and metabolic pathway-based analyses indicate that calculus preserves dental plaque biofilms that calcify at different stages of biofilm development, which does not directly reflect any of the oral pathologies that we have recorded.

### Comparison with pre-tobacco introduction populations in Europe

While we did not detect differences in the microbial species profiles or microbial metabolic pathway profiles between heavy smokers and light/non-smokers within the MID and CMB individuals, these populations were living during a time when smoking was common and many people would have been exposed to high levels of second-hand smoke, even if they were not using pipes or smoking themselves. High second-hand smoke exposure might obscure smoking-related species profile changes that develop between smokers and non-smokers (Beghini et al., 2019). To investigate possible differences in species profiles that are related to smoke exposure, we chose to broadly compare calculus species profiles of European populations living before and after the introduction of tobacco to Europe during three time periods: Medieval, Industrial, and present-day Modern.

We selected a total of 6 additional European populations based on geographic proximity and availability of comparative samples (Figure 5A). As pre-tobacco Medieval populations, we included the Kilteasheen calculus data set (KIL) (Mann et al., 2018) from Ireland. Additionally, we produced data from two sites in Spain dated to the medieval period, El Raval (ELR) and Iglesia de la Virgen de la Estrella (IVE), to match the CMB population. As an additional Industrial-era smoke-exposed population, we included the Radcliffe calculus data set (RAD) (Velsko et al., 2019), from the early 1800s England. Modern calculus data sets from Jaen, Spain (JAE) (Velsko et al., 2019) and from Valencia, Spain (VLC) (Fellows Yates et al., 2021b) were also included for comparison. Poorly preserved samples were removed based on preservation assessments (Supplemental figures S2, S3), leaving 176 samples that were used in downstream analyses (MID = 65, CMB = 8, KIL = 35, ELR = 3, IVE = 5, RAD = 42, JAE = 10, VLC = 8).

We first wanted to know if there is a change in the average number of species detected between samples from the Medieval, Industrial, and Modern periods. There are statistically significant differences between Modern and Medieval groups (Figure 4, p < 0.001, effect size 0.71), and between Modern and Industrial groups (Figure 4, p < 0.001, effect size 0.48), but also between Industrial and Medieval groups, although the effect size was small (p < 0.01, effect size 0.24). We found few differences between historic sites, with significant differences between only MID and CMB (p < 0.001, effect size 0.37) and between MID and KIL (p < 0.001, effect size 0.41). All historic sites had significantly fewer species than either of the modern groups, VLC or JAE (Supplemental figure S6, Supplemental Table S9). The Shannon index was not significantly different between any time periods (Figure 4B) or sites (Supplemental figure S6), indicating that the distribution of species is highly similar across all samples.

**Figure 4.**
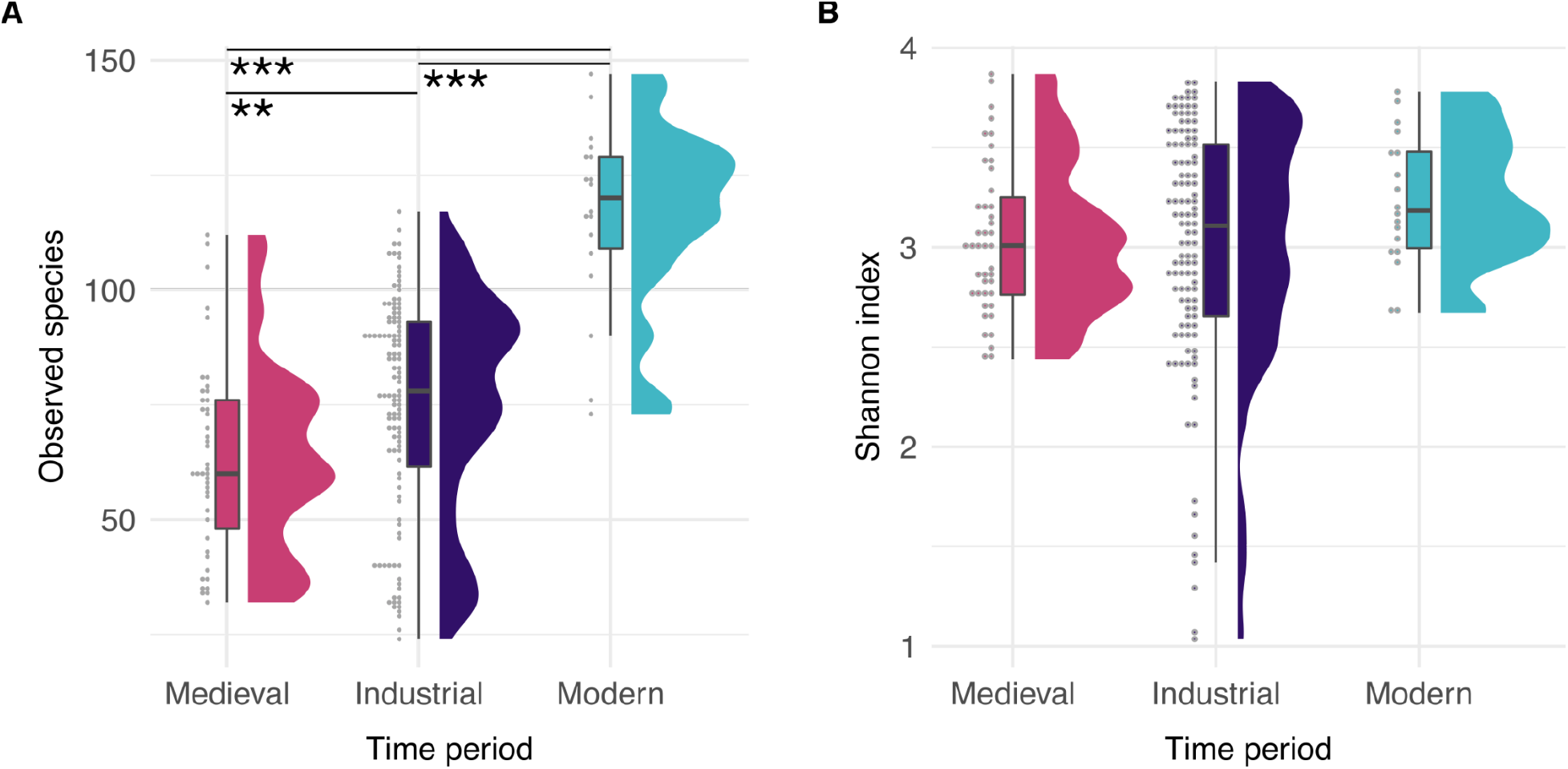
Within-sample species diversity. **A.** Raincloud plots showing the observed species in each sample, grouped by time period. **B.** Raincloud plots showing the Shannon index in each sample, grouped by time period. *** p < 0.001, ** p < 0.01.

To confirm the trend of increasing numbers of species in calculus samples over time, we investigated the influence of sequencing depth and average read length on species detection. The modern calculus samples were sequenced much more deeply than the historic samples, and they have a longer average read length, possibly making detection of low-abundance taxa more likely. We generated two additional datasets by down-sampling all calculus libraries in two ways: first by sequencing depth, then by read length. For the first set, we randomly subsampled all libraries with > 10M reads down to 10M reads, while maintaining all libraries with < 10M reads (Sub 10M set). For the second set, we subsampled all libraries to include only reads ≤ 75bp in length (Sub 75bp set). Each subsetted dataset was then profiled with MetaPhlan3, and the number of species and Shannon index calculated (Supplemental Figures S7, S8).

Both subsampling methods reduced the number of species detected for all groups (Supplemental Figures S7A, C, and S8A, C), however Modern samples still had significantly more species detected than either Medieval or Industrial samples. The Shannon index was unaffected by subsampling for read depth (Supplemental Figures S7B,D), but was affected by subsampling for read length (Supplemental Figures S8B,D). We found that the Sub 10M dataset had significantly fewer reads than the full set for most sites (Supplemental Figure S9A), but the average read length and the average GC content of the reads were unaffected (Supplemental Figure S9B,C). The Sub 75bp dataset compared to the full dataset had significantly fewer reads (Supplemental Figure S9A) and significantly lower average read length (Supplemental Figure S9B) for most sites, but the average GC content was minimally affected (Supplemental Figure S9C). The number of species detected in the Sub10M dataset compared to the full dataset was significantly lower for both the Modern sites but none of the historic sites, while the number of species detected in the Sub75bp dataset compared to the full set was significantly lower for all sites, although two were not significantly different (Supplemental Figure S10). The subsampling results suggest that the higher species counts in Modern samples compared to Medieval and Industrial samples may be a real effect, but further detailed investigation of normalizing sequencing depth and read lengths, and addition of more modern calculus samples from more sites, are needed to confirm this.

We next wanted to determine if the microbial community structure of calculus from Medieval, Industrial, and Modern calculus groups differ. This would let us know whether there are substantial changes that have occurred in species composition of dental calculus between distinct historic periods. We performed PCA based on the species abundance table to see whether the samples clustered by time period or by other metadata categories (Figure 5B, Supplemental Figure S11A). The Medieval samples generally cluster away from the Industrial samples along PC2, while the Modern samples generally cluster with Industrial and Medieval samples at one end of PC1. While this may suggest that there are time-related differences in species composition driving separation of samples, we note several other factors that may be driving the species composition pattern and confounding this observation.

**Figure 5.**
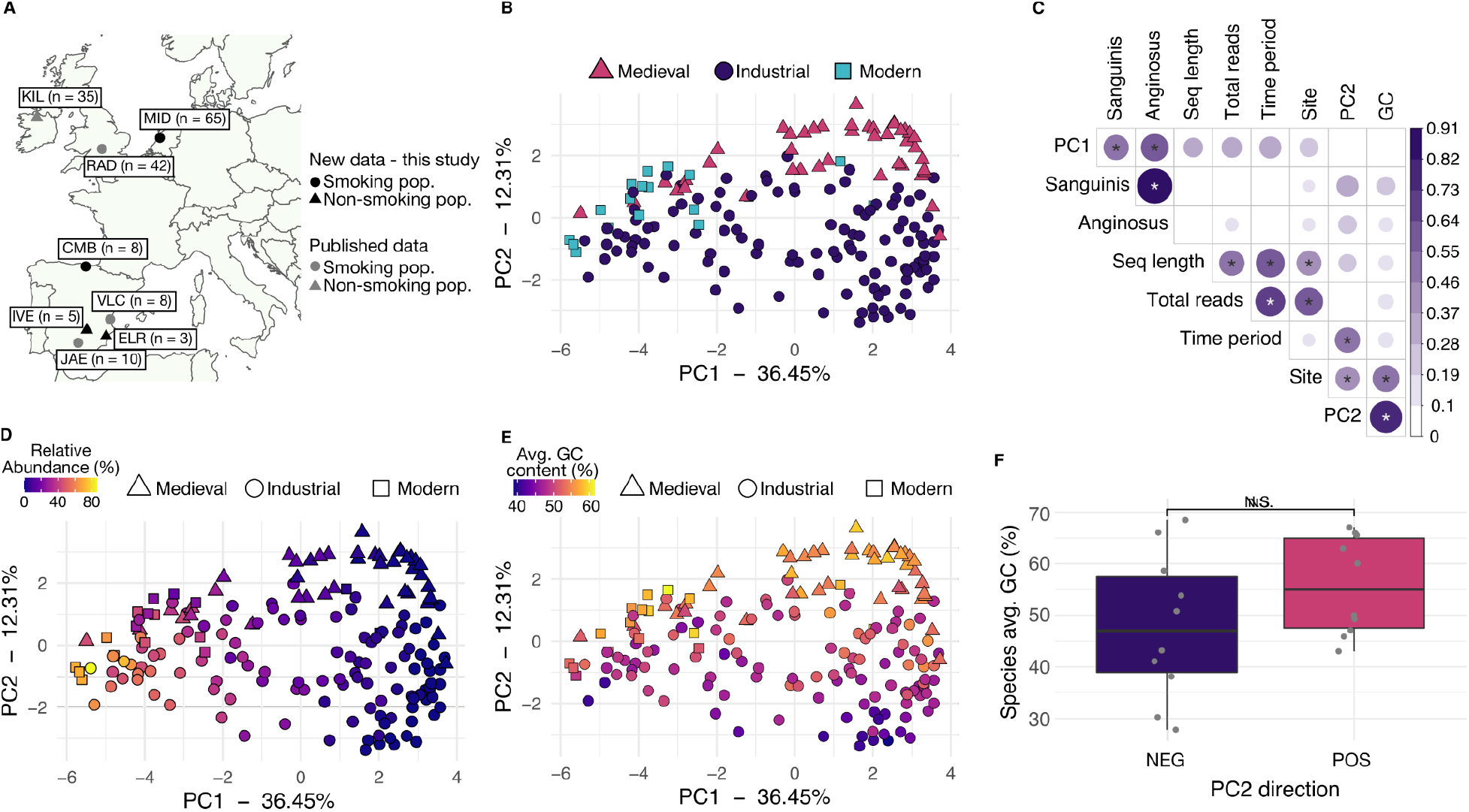
Community structure is shaped by species aerotolerance and sample GC content rather than time period. **A.** Map of sample sites. **B.** PCA of species profile colored by time period. **C.** Canonical correlation (CC) analysis correlations between metadata categories principal component loadings for all libraries. Significance tests were performed with a Pearson correlation test. The size and color of the dots corresponds to the CC value, which does not determine the direction of the correlation (positive or negative. Hence all CC values are positive). The tested metadata were selected because the information was available for the majority of libraries. Only correlations ≥ 0.4 have significance indicated with stars, * p ≤ 0.001. **D.** PCA of species profile colored by relative abundance of the 10 species with strongest negative loadings in PC1, all of which are aerobic or facultative species found early in dental biofilm development (Supplemental Table S2). **E.** PCA of species profile colored by average GC content of the sample. **F.** Average GC content of the 10 species with strongest PC1 negative (NEG) and positive (POS) loadings, indicating that species characterizing the samples with higher average GC content have higher average GC content than the species characterizing the samples with lower GC content. N.S - non-significant (p > 0.05 by Wilcox test). **PC1** - PC1 loadings; **Sanguinis** - proportion of total reads that were assigned to a species in the Sanguinis group; **Anginosus** - proportion of total reads that were assigned to a species in the Anginosus group; **Seq length** - library average sequence length; **Total reads** - total reads in the library after quality-trimming and merging; **Time period** - time period of the samples; **Site** - site of the samples; **PC2** - PC2 loadings; **GC** - library average GC content.

Canonical correlation analysis revealed there are significant correlations between PC1 loadings and the proportion of Sanguinis group and Anginosus group *Streptococcus* present in samples, and there are significant correlations between PC2 loadings and library average GC content, in addition to site and time period (Figure 5C). We further investigated the sources of variation in the samples that most strongly influence PC loadings. Samples appear to separate along PC1 according to biofilm maturation stage (Figure 5D, Supplemental Figure S11B), similar to Figure 2, however the loadings have reversed the species gradient. The samples with more negative PC1 loadings, including all but one of the Modern calculus samples, are enriched in early colonizer, aerobic and facultative taxa such as *Streptococcus*, *Neisseria*, and *Rothia* (Supplemental Table S4). Modern calculus samples are known to have higher levels of early-colonizer species than historic European dental calculus, which may be related to toothbrushing and other dental interventions (Velsko 2019). In contrast, the samples with more positive PC1 loadings are enriched in late colonizer, anaerobic, proteolytic taxa such as *Treponema*, *Tannerella*, and *Fretibacterium* (Figure 5D, Supplemental Figure S11B, Supplemental Table S4). The late-stage colonizer *Methanobrevibacter oralis* is highly abundant in several Industrial-era samples and we performed an additional PCA on a table without *M. oralis* to check if this species was strongly shaping the PCA, and found that it is not (Supplemental figure S12).

Loadings in PC2 are correlated with GC content, time period, and site (Figure 5C), which are themselves correlated. Only a single time period is represented per site, and older calculus samples are expected to have higher average GC content due to the taphonomic loss of short AT-rich fragments (Fagernäs et al., 2020; Mann et al., 2018). Thus it is expected that the Medieval calculus samples have higher average GC content than the Industrial samples. This is, however, not entirely an age-related effect, as the modern samples also have a higher average GC content, in the same range as the medieval samples (Figure 5E). The species characterizing samples with high positive PC2 loadings have a higher average GC content than the species characterizing samples with strong negative PC2 loadings (Figure 5F), indicating the trend in GC content is reflected in taxonomic assignments. The average read length of samples does not appear to influence the species composition of these samples, as the modern samples and Radcliffe samples have the longest read lengths and are distributed across PC1 from the lowest to the highest values (Supplemental Figure S7C). Taken together, it appears that the factors driving differences in species diversity between samples are derived from a variety of sources, several of which are unrelated to biofilm and host ecology. These results indicate human behavioral differences such as tobacco smoking may not be as strongly reflected in calculus community profiles as they are in dental plaque, and studies will need to be carefully constructed to account for this and maximize the chance of observing signals related to the study question.

### Streptococcus species distributions

Although we did not find associations between oral pathology metadata and species profiles of MID calculus samples, we found differences related to species ecological niches. This suggested an opportunity to investigate distinctive distributions of *Streptococcus* species in our dataset, such as that reported in a deep-time study of ancient calculus (Fellows Yates et al., 2021b). Fellows Yates, *et al*, reported a small group of human calculus samples that had a low proportion of *Streptococcus* that was represented predominantly by species of the Anginosus group, rather than of the more typical Sanguinis group, which is a pattern that resembles streptoccocal patterns found in chimpanzee calculus. Fellows-Yates et al. (2021b) were unable to speculate on the reasons for this distinction within humans because of small sample size and insufficient metadata, but the MID collection presents the opportunity to examine a large number of dental calculus samples from a homogenous population sampled from a single location and time period with associated high quality health-related metadata.

To determine whether the distribution of *Streptococcus* groups across humans is consistent in our populations, or whether there are humans with a *Streptococcus* group profile that more closely resembles that found among chimpanzees as reported by Fellows Yates, *et al.* (Fellows Yates et al., 2021b), we used a species table generated by MALT using the same RefSeq database used by Fellows Yates, *et al.* This allowed us to directly compare our results with previously published results. We grouped all *Streptococcus* species in each individual into seven phylogenetically-supported clades (Richards et al., 2014): Sanguinis, Mitis, Salivarius, Anginosus, Bovis, Pyogenic, and Mutans, as well as the categories Other if they fell outside of these clades, or Unknown if they have not been phylogenetically placed.

When the individuals from MID and CMB were ordered by decreasing proportion of Sanguinis group *Streptococcus*, there was a clear distinction within the population, with nine of 72 individuals (11%) having Anginosus as their dominant group (> 50% of all *Streptococcus* reads were assigned to species in the Anginosus group), as opposed to Sanguinis for the majority of individuals (Figure 6A). These Anginosus-dominated individuals had a lower overall proportion of *Streptococcus* (Figure 6B), which was due to a loss of Sanguinis group species (Figure 6C), and were the same individuals that have species profiles characterized by a high proportion of anaerobic taxa (Supplemental Figure S5).

**Figure 6.**
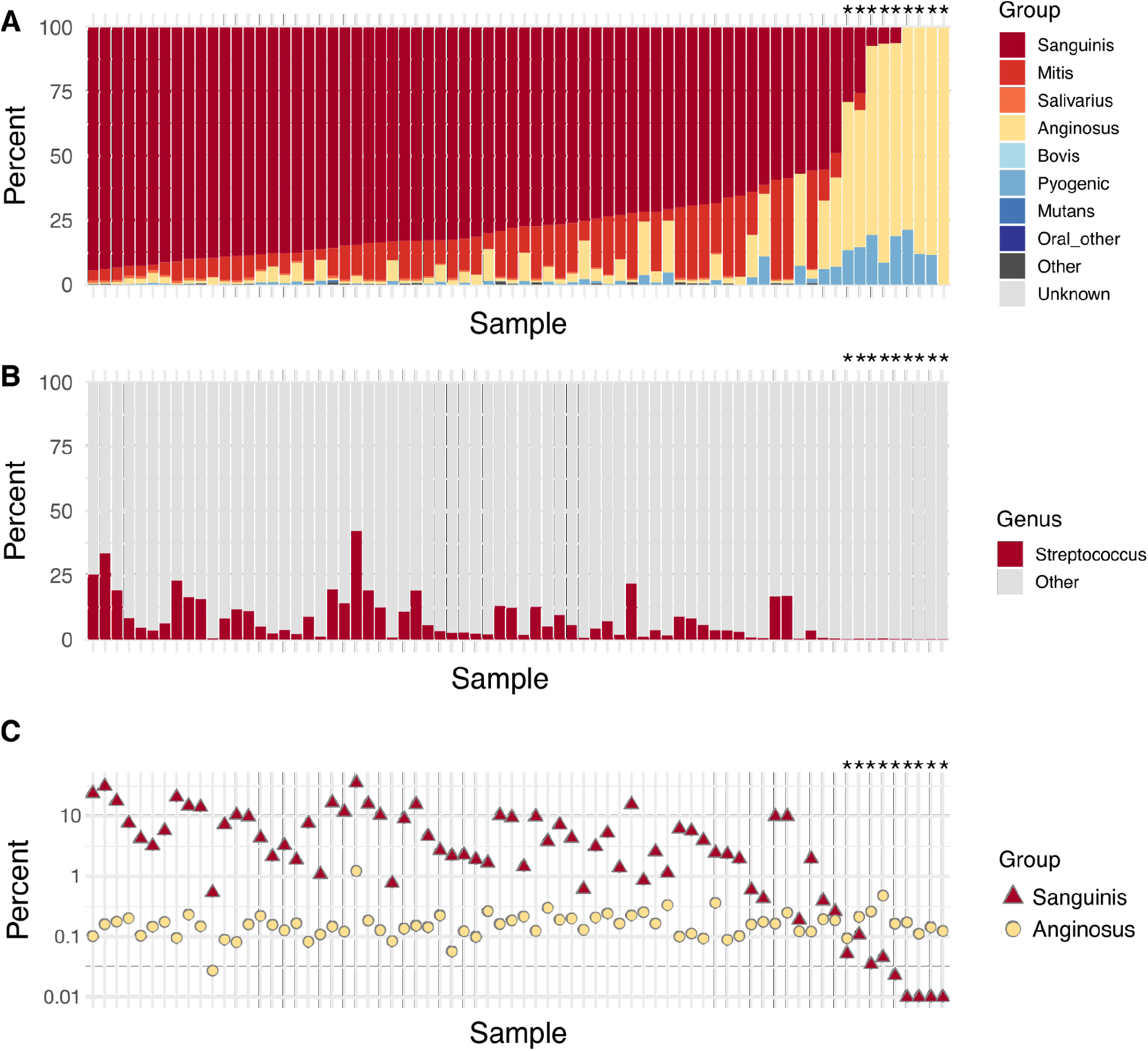
*Streptococcus* species group distributions in Middenbeemster and Convento de los Mercedarios de Burtzeña. **A.** Percent of *Streptococcus*-assigned reads in each *Streptococcus* group per sample, ordered by decreasing proportion of Sanguinis group and increasing proportion of Anginosus group. **B.** Percent of reads assigned to *Streptococcus* out of all genera detected per sample. **C.** Percent of reads assigned to species in the Sanguinis and Anginosus groups per sample. The y-axis is log-scaled and the four samples on the far right have no reads assigned to Sanguinis group *Streptococcus* species. Sample order is identical across all 3 panels. A star (*) indicates samples in which >50% of the *Streptococcus* reads come from species in the Anginosus group.

Since this pattern of *Streptococcus* species distribution has now been shown at a continental scale (Fellows Yates et al., 2021b) and within two individual populations (MID, CMB), we next asked whether this pattern holds for an individual, or whether there is intra-individual variation. To investigate tooth-specific *Streptococcus* species differences with individuals, we used the data published by Fagernäs, et al. (Fagernäs et al., 2022). This dataset consists of calculus sampled from 12-18 teeth per individual from 4 individuals. The species table published by Fagernäs, et al. was generated with the same version of MALT and the same RefSeq database, and we therefore grouped the *Streptococcus* species as described above, per tooth. When multiple sites of a single tooth were sampled, we averaged the read counts to produce a tooth-wide average.

We found that three of the individuals (CM55, CM59, and CM82) had a majority of teeth dominated by Sanguinis group *Streptococcus* species (Figure 7A,B, Supplemental Figure S13), while in individual CM165, 6 of 15 teeth (40%) were dominated by Anginosus group *Streptococcus* species. Individual CM165 had overall, across their entire dentition, a lower proportion of reads identified as *Streptococcus* compared to the other three individuals (Figure 7C), where those teeth with the lowest proportion of *Streptococcus* reads were dominated by Anginosus group species (Figure 7D, Supplemental Figure S13D). Across each individual in this data set, teeth with lower proportions of *Streptococcus* had lower proportions of Sanguinis group species and higher proportions of Anginosus group species, indicating that this dichotomy is a tooth-specific phenomenon that is reflected at an individual level.

**Figure 7.**
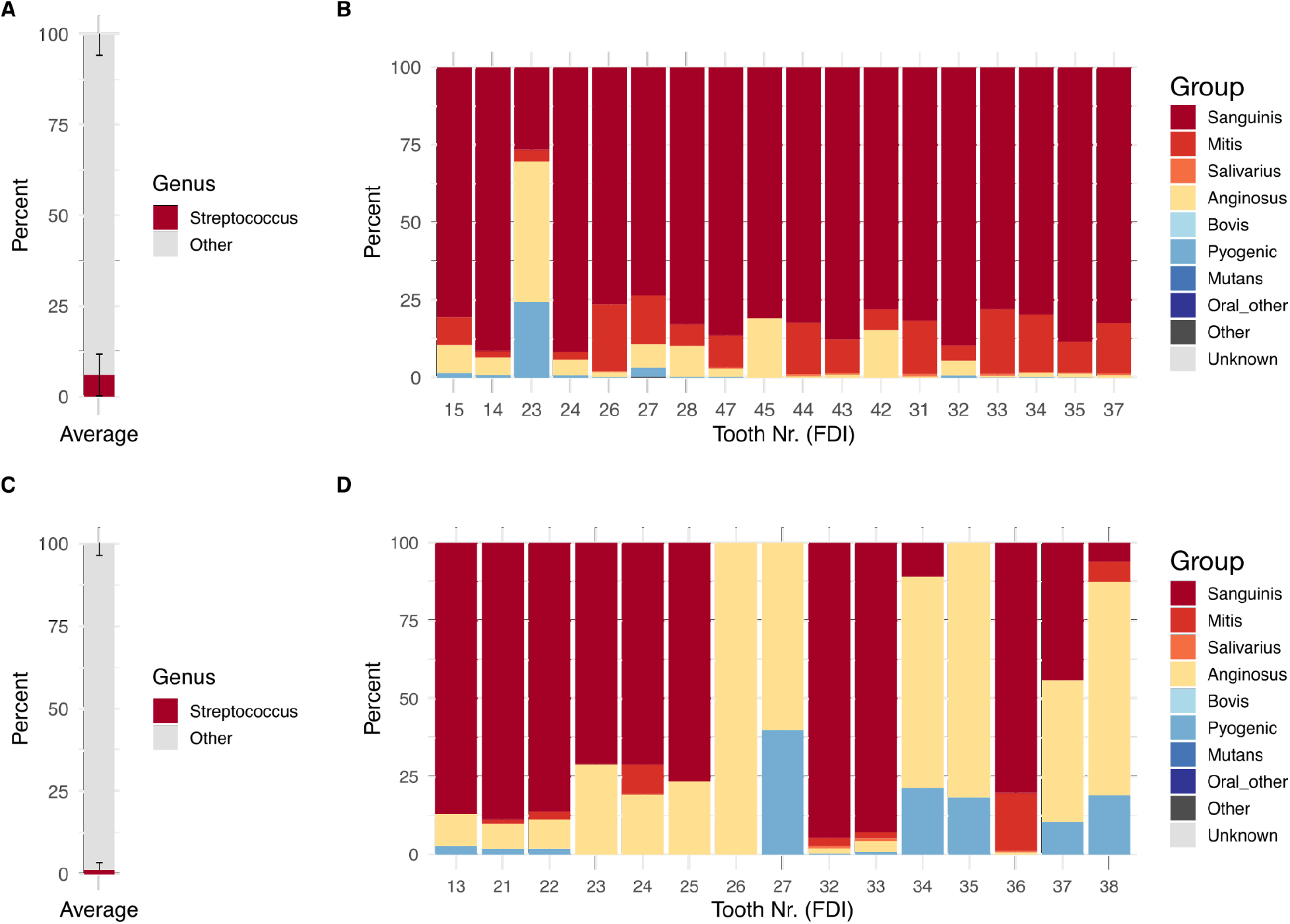
Distribution of *Streptococcus* groups in calculus of each tooth sampled from two individuals from the Chalcolithic site (ca. 4500-5000 BP) Camino del Molino, Spain. **A.** Average proportion of reads assigned to the genus *Streptococcus* compared to all other genera in individual CM55, averaged across all teeth sampled. **B.** Proportion of reads assigned to species within each group of *Streptococcus*, by tooth, in individual CM55. **C.** Average proportion of reads assigned to the genus *Streptococcus* compared to all other genera in individual CM165, averaged across all teeth sampled. **D.** Proportion of reads assigned to species within each group of *Streptococcus*, by tooth, in individual CM165. Tooth numbers are in FDI World Dental Federation notation.

## Discussion

We investigated patterns related to dental health and pipe use in ancient dental calculus in a large osteological collection from the Netherlands, as well as a small group from Spain. No microbial species distribution patterns pertaining to dental health were detected in the samples, nor was there any indication that species profiles distinguish individuals with pipe notches (heavy smokers) and those without pipe notches (light/non-smokers). Further, a broader time scale, spanning the medieval period through modern day, also did not show patterns of differentiation based pre- and post-introduction of tobacco to Europe. Instead, patterns appear to be driven by individual variation in biofilm development, with individuals from each time period having species profiles ranging from more aerotolerant to more anaerobic. This pattern is highlighted in the proportions of *Streptococcus* groups detected in calculus, where a minority of individuals carry exceptionally low proportions of Sanguinis group *Streptococcus* species, appearing instead to be dominated by Anginosus group species. However, this minority has overall very low proportions of *Streptococcus* species, and high proportions of late-colonizer anaerobic taxa, and this pattern appears to be an individual-specific trait.

Although there were some differences in dental pathologies by pipe use status within the Middenbeemster collection, with users having fewer caries, more severe periodontal disease and larger calculus deposits, it was difficult to correlate the results directly with pipe smoking for two key reasons. First, the clear relationship between pipe use and biological sex, with the majority of males having at least one pipe notch and most females having no pipe notches, confounded comparison of pipe users and non-users. In many populations, including Dutch populations of the period (Baetsen and Weterings-Korthorst, 2013; Clevis and Constandse-Westermann, 1992; Jackes, 2015), there are significant differences in oral health between males and females that relate to biological differences and gendered differences in diet (Lukacs and Largaespada, 2006; Lukacs, 2008). Without having more information from female pipe users and male non-pipe users it is difficult to disentangle whether the differences we see are related to biological or gender differences.

Another issue is that there is a strong correlation between age and the presence and severity of oral pathology, and because our sample was drawn from a natural cemetery assemblage, it was not balanced in terms of age: the smokers/males were older. A pertinent issue here is inability to know whether individuals with significant tooth loss were smoking pipes. However, a previous study comparing oral pathology between Middenbeemster males and earlier pre-tobacco males from Klaaskinderkerke (Figure 1A) identified lower caries rates, and higher AMTL and calculus severity in pipe smoking males in comparison to pre-tobacco males (Inskip et al., n.d.). Similar results were also found by others who had much larger sample sizes with a better representation of pipe users and non–users (Geber and Murphy, 2018; Walker and Henderson, 2010; Western and Bekvalac, 2020). While these results suggest an impact, a confounding issue is that it is not possible to entirely disentangle the influence of smoking versus the abrasion of teeth by clay pipe stems, which may lead to the obliteration of early caries, hence lower caries rates, but also speed up tooth loss due to advanced wear.

The Middenbeemster skeletal collection is an extensively studied assemblage, and a wealth of metadata is available for each individual. Given that dental plaque biofilm species profiles are distinct between healthy and disease-affected teeth (Abusleme et al., 2021), studying this collection afforded us the opportunity to assess whether ancient dental calculus microbial communities also maintain differences related to dental health. While we found no broad community-scale differences relating to pathology including periodontal disease, calculus score, caries, and antemortem tooth loss, among others, this is consistent with the observation that dental calculus appears to be a stable climax biofilm community (Velsko et al., 2019). The ecological succession by which biofilms reach a terminal state (Kolenbrander et al., 2006; Listgarten et al., 1975, 1973; Socransky et al., 1977; Theilade et al., 1974, 1966; Tinanoff et al., 1976) may be influenced by external forces including oral hygiene, immunological responses, and tobacco use, with intermediate biofilm states strongly depending on the local oral environment (Wade, 2021). However, once a terminal community is reached, the fully mature biofilm may have lost most indications of earlier community states.

Tobacco smoking adversely affects oral health, and is associated with substantial changes in dental plaque biofilm community profiles (Buduneli, 2021; Nociti et al., 2015). Whether the microbial changes are due to direct influence of tobacco smoke exposure or to physiological changes in the oral tissues is not clear, and different studies report different specific species abundance changes (Al Bataineh et al., 2020; Kumar et al., 2011; Mason et al., 2015; Moon et al., 2015; Yang et al., 2019). Overall, however, the species profile appears to shift to one with more pathogenic potential, with elevated abundances of anaerobic, proteolytic species that are associated with periodontal disease. Given that dental calculus species profiles generally have higher abundance of these same disease-associated species, even in the absence of dental pathology (Kazarina et al., 2021; Velsko et al., 2019), detecting smoke exposure-induced changes may be difficult. Specific changes might be detected by differential abundance analysis, but the relative value of this information in ancient dental calculus remains to be investigated.

The patterns we detected within the MID and CMB samples appear to be related to individual differences in the terminal dental biofilm community. The results of our transect support this observation, which is consistent with other publications that have performed time transect studies (Fellows Yates et al., 2021b; Ottoni et al., 2021), where community diversity analyses do not cluster samples by time period. Instead of patterns relating to sample site, time period, or dental health, the samples appear to maintain individuality. This pattern of stable microbial signatures in dental plaque is a known phenomenon, with individual teeth having relatively stable communities over months-long time scales (Tamashiro et al., 2021). Despite high variability in profiles across teeth within an individual, plaque from any one tooth in an individual is more similar to other teeth in the same individual than to teeth in other individuals (Tamashiro et al., 2021). This pattern of individuality is also captured in ancient dental calculus (Fagernäs et al., 2022), where there is relatively high variability across teeth in an individual, yet samples from one individual are more similar to each other than to samples from another individual.

The strongest pattern we detected separating samples in beta-diversity analysis was the proportion of species with different environmental niches. This was shown by a strong gradient where samples at one end of the PCA were enriched in aerotolerant, early biofilm colonizer taxa, and samples at the opposite end were enriched in anaerobic, later biofilm colonizer taxa. This was true for not only the MID and CMB samples, but also for samples from other sites and time periods in Europe. We found no correlation with dental pathology, suggesting individual differences in plaque development timelines may account for this pattern, and studies of biofilm succession may offer insight. *In vivo* studies of plaque development have reported two distinct patterns, one termed “rapid” and the other “slow” (Listgarten et al., 1975; Zee et al., 1997, 1996). Slow plaque-forming individuals maintained a “younger” biofilm dominated by more aerotolerant taxa, including *Streptococcus* and other Gram-positive cocci (Listgarten et al., 1975; Zee et al., 1997, 1996). During the same period of time, the biofilm in rapid plaque formers developed a more complex community with higher proportions of anaerobic and Gram-negative taxa. This difference in the rate at which biofilms mature and calcify, and the species composition of the end-state climax community, may be captured in dental calculus.

We found the abundance of *Streptococcus* and the distribution of species within distinct phylogenetic lineages of *Streptococcus* to reflect this individual difference. While Fellows Yates, *et al.* (2021b) identified a small number of human calculus samples that have a *Streptococcus* species profile and abundance distribution that is more similar to chimpanzee calculus, they were unable to further speculate on the origin of this distinction. We found this pattern replicated in the MID and CMB samples, and with no association with dental pathology. Finer-scale assessment of four individuals for which data from multiple calculus samples was available from across their dentition revealed that the abundance and distribution of *Streptococcus* species was fairly uniform in three individuals but variable in the fourth. This fourth individual had variable amounts of *Streptococcus* across their dentition and the predominant species were distinct between teeth with relatively high and relatively low *Streptococcus* abundance. Overall this individual had fewer reads assigned to *Streptococcus*, and fell within the ‘chimpanzee-like’ profile.

The results of this study suggest that it may not be possible to use dental calculus metagenomes to distinguish broad health-associated changes in community profiles such as can be detected in living dental plaque biofilms. Instead, the value of these ancient metagenomes may lie in providing insight into individual biofilm characteristics, particularly related to biofilm and microbial ecology. Another intriguing avenue of ancient dental calculus metagenome research is on the evolution of specific species and strains through the assembly and reconstruction of ancient metagenome assembled genomes (Brealey et al., 2020; Granehäll et al., 2021; Wibowo et al., 2021). In contrast, the wealth of proteins and metabolites that are preserved in calculus may reflect biofilm community responses to altered oral environments such as dental disease or tobacco smoking (Jersie-Christensen et al., 2018; Velsko et al., 2019, 2017), and could be used instead of the metagenome to study the role of health in shaping the oral microbiome in deep time.

## Materials and Methods

### Archaeological sites and associated skeletal remains

#### Middenbeemster (MID)

Middenbeemster is today a small town in the Beemster region of the Dutch province of North Holland (Figure 1A). Settlement in the region began after the draining of the Beemster lake in 1612 (Aten et al. 2012). Following land reclamation, the region was used primarily for agriculture, and it became particularly renowned for its dairy farming and cheese production (de Jong et al., 1988), although residents also worked in local service industries, shops, and the military (Falger et al., 2012). While it was initially planned to construct multiple churches throughout the region, only one church, the Keyserkerk, was completed at Middenbeemster, and as such it served the entire Beemster community (Aten et al., 2012).

In 2011, the Laboratory for Human Osteoarchaeology of Leiden University and the company Hollandia Archeologen carried out archaeological excavations at the Middenbeemster Keyserkerk prior to a planned construction project. The excavation of part of the cemetery (Leroux, 2012), which was in use from 1623 to 1866, uncovered approximately 450 individuals, and their skeletal remains were relocated to the Human Osteology Laboratory of Leiden University for research and long-term care. Most of the excavated individuals date to the later periods of the cemetery’s use because historical records indicate that the older portions of the cemetery, which had become overcrowded, were cleared in 1829 to make room for new burials (Leroux, 2012). At the time of death, undertakers and gravediggers wrote down individual information for each burial, including name, age at death, date of death, and the profession of the deceased, creating burial records, death registers, and a map of Middenbeemster’s cemetery (Veldman, 2013). This information was used to identify some of the excavated skeletons; however, alternative ways of spelling names and typographical errors have made it difficult to identify most remains with certainty (Leroux, 2012). After excavation, the skeletal remains recovered from the cemetery were washed with water to remove clay and sand from the burial prior to storage.

All metadata were collected by the Laboratory for Human Osteoarchaeology of Leiden University and confirmed by S. I. and M. H.. Biological sex estimates were made by observing and recording the sexually dimorphic morphological traits of the skull and pelvis, following standard methods (Buikstra and Ubelaker, 1995; Workshop of European Anthropologists, 1980). The approximate age at death for each individual was estimated using dental attrition (Maat, 2001), cranial suture closure (Lovejoy et al., 1985), the sternal rib-end (Işcan and Loth, 1986), the pubic symphysis (Brooks and Suchey, 1990), and the auricular surface (Buckberry and Chamberlain, 2002) and grouped into four age categories: adolescent (13-18 years), young adult (18-25 years), middle adult (25-45 years), and old adult (>45 years). For known individuals, age estimates were cross checked with archival information, which showed a high degree of accuracy for sex estimates, and good accuracy with age estimates.

Total DNA from the calculus of 75 Beemster individuals was extracted and sequenced for this study. The oral health of each skeleton was recorded with reference to the presence and absence of pipe notches, periapical lesions, supra- and subgingival calculus scores, periodontal diseases, caries, antemortem tooth loss, and their respective severeness (Supplementary Table S1). To be scored in the analysis of dental pathology, it was important to know whether an individual had pipe notches or not. As such, only individuals retaining at least 50% of their anterior teeth in alternating positions (thereby allowing pipe notches to be observed) were included in the analysis of dental pathology. To evaluate the presence and severity of periodontal disease, the mandible and maxilla of each individual were examined macroscopically. Changes were recorded based on the inflammatory loss of the alveolar bone, and scored following the four point scale described in (Ogden, 2007): None – 1, Mild – 2, Moderate – 3, Severe – 4. Additionally, the percentage of tooth positions with signs of periodontitis was calculated.

Caries were recorded when there was evidence for enamel loss (Hillson, 2001) on any tooth surface; this included the lingual, buccal, occlusal, medial, and distal crown sites, as well as at the cementoenamel junction (CEJ) and root surfaces. Gross caries were scored when it was no longer possible to identify the origin of the carious lesion. Individuals were recorded as having caries if one or more teeth were affected and caries rates per individual were also calculated. Antemortem tooth loss was scored as present when one or more teeth were absent and there was evidence of remodeling to the alveolar margins of the socket that indicated healing. The total number of teeth lost antemortem was calculated. Calculus was graded following the four point scale by Brothwell (Brothwell, 1981): 0=none, 1=slight, 2=moderate, 3=extensive. Due to the complexity of differentiating between types of periapical lesions (granuloma, abscess or cysts) without x-ray, an individual was scored as positive for the conditions when at least one of the three types, as identified by Ogden (2008), were present. For periodontal disease and calculus, individuals were given a score based on the highest observed score per individual. A full table of the collected dental metadata is in Supplementary Data S1.

Dental calculus from the Middenbeemster collection was collected by K. Z. in 2014. For each individual, calculus from multiple teeth was sampled and pooled for analysis. Calculus from a total of 75 individuals was sampled. Of these, 40 individuals had pipe notches, and 7 individuals had dentitions that were too fragmentary to determine whether pipe notches were present.

#### Convento de los Mercedarios de Burtzeña (CMB)

The Convento de los Mercedarios de Burtzeña, is located in the eastern part of Burtzeña, in Northern Spain. Archaeological and historical information about the site and its excavation are available in (Domínguez Ballesteros et al., 2018) and (García-Collado et al., 2018). The use of the cemetery is dated between the end of the 16th century. and the beginning of the 19th century. The excavations revealed the remains of the foundation under the pavement of the church, which contained the sandstone-boxed inhumations of more than 150 graves, of which most were occupied by several individuals. Fifty graves, covered with slabs, were excavated during the excavation works. A total of 62 articulated individuals and numerous scattered bone remains were found. Each grave contained individuals of different ages, which may represent family graves, which were typical at that time.

All metadata for the site were investigated and collected by M. I. G-C., E. D. B. and L. S. Z.. All the anthropological remains recovered were cleaned by washing with water at room temperature and brushing with soft bristled brushes. Care was taken not to submerge the bones completely in order to minimize water absorption. The material was then left to dry at room temperature on absorbent paper in a ventilated room.

Calculus from 8 individuals from the Convento de los Mercedarios de Burtzeña cemetery were extracted and sequenced for this study. Four individuals had pipe notches in their dentition, and four did not. It is possible that individuals CMB003 and CMB004 are the same individual, as each is represented by a partial mandible recovered from the same grave. The oral health of each skeleton was recorded with reference to the presence and absence of pipe notches. This was investigated as previously described (García-Collado et al., 2018). Biological sex was estimated observing the sexially dimorphic morphological traits of the skull and pelvis (Buikstra and Ubelaker, 1995; Phenice, 1969). Age-at-death of each individual was estimated using the pubic symphysis (Brooks and Suchey, 1990), the auricular surface (Lovejoy et al., 1985), the sternal rib-end (Işcan and Loth, 1986) and the sacrum (Passalacqua, 2009). Since most individuals originated from disarticulated assemblages of human remains and often were just isolated mandibles, it was not possible to make precise age estimations for all skeletons. Four individuals could only be classified as “older than 20 years” and therefore could not be classified into the same age groups as used for the Middenbeemster collection; they were instead classified as “Adult”.

Dental calculus from the Convento de los Mercedarios de Burtzeña was collected by M. I. G-C. in 2019. For each individual, calculus from multiple teeth was sampled and pooled for analysis. Eight individuals were sampled, four with pipe notches, and four without pipe notches (Supplemental Figure S1).

#### El Raval and Iglesia de la Virgen de la Estrella

Dental calculus was also analyzed for two Spanish sites dated to the Medieval period, El Raval (ELR) and Iglesia de la Virgen de la Estrella (IVE). Calculus from five individuals from each site was extracted and sequenced for this study (Supplemental Table S1). Samples from El Raval were collected in 2018 by D.C.S.G. Calculus were collected off of multiple teeth and pooled for extraction/collected off a single tooth per individual (Salazar-García et al., 2014).

El Raval is a medieval necropolis from the city of Crevillent (Alacant, Spain), from the times of the Kingdom of València, Crown of Aragón. The necropolis was radiocarbon dated between the end of the 14^th^ century AD and the beginning of the 16^th^ century A.D., and was located outside the city wall besides one of the main roads leading into the city (Martí et al., 2009). Burial customs revealed that most individuals interred were Mudéjar, Muslims of Al-Andalus that remained in Iberia after the Christian Conquest, and a minority were Islamic people who had converted to Christianity (Trelis et al., 2010). A total of 81 burials were recovered, mostly single graves occasionally covered by rocks or wood. Individuals of all ages, except those older than 60, were buried in the cemetery, being both full adults and infants (0-4 yo) the most frequent. Oral pathologies, such as dental calculus, caries, periodontal disease and antemortem tooth loss, are frequent amongst the adult individuals (de Miguel Ibáñez, 2007).

Samples from Iglesia de la Virgen de la Estrella were collected in 2016 by D.C.S.G. Calculus were collected from multiple teeth and pooled for extraction.

Metadata for the samples sequenced for this study are in Supplemental Table S1.

### Comparative data sets

Two published datasets were selected to represent comparative European populations pre- and post-introduction of tobacco to Europe: the Kilteasheen (KIL) dataset from Ireland ca. 600– 1300 CE (n = 36, (Mann et al., 2018)), and the Radcliffe Infirmary (RAD) burial ground set from England ca. 1850 (n = 44, (Velsko et al., 2019)). Data from Chalcolithic-era dental calculus (ca. 4500–5000 BP) previously reported in Fagernäs, et al. (Fagernäs et al., 2022) were included as a comparative dataset to examine intraindividual variation in *Streptococcus* species. The samples are listed in Supplemental Table S5.

### Tobacco use at the sites

European encounters with tobacco commenced in the15th and 16^th^ century during the European colonization of the Americas, at which time it was presented to them by indigenous American peoples (Gately, 2001; Norton, 2008). Tobacco was long used by indigenous peoples in a myriad of ways; it formed an important part of their lives and identities. Its use as a medicinal agent, inspired Europeans to investigate its healing and curative properties, while its role in ritual and political ceremonies demonstrated its use as a social and recreational entity (Norton, 2008). In terms of smoking, the habit traversed the Atlantic via colonialists, returning adventurers and sailors, sojourners, Indigenous delegates, and enslaved individuals. From there it diffused into the general population (Brongers, 1964; Goodman, 1993; Norton, 2008). By the early 17^th^ century, tobacco was a taxable commoditythorughout much of western Europe including present-day Spain, England, France, and the Netherlands, and was consumed by a large proportion of society (Goodman, 1993).

Tobacco smoking is well documented in the Netherlands during the 17th to 19th century. Initially, the Dutch procured much of their tobacco from the English, although later they had their own domestic industry (Brongers, 1964). While there were changing fashions, clay pipe smoking was the dominant method for tobacco use in the Netherlands, who became the leading producers of pipes in Europe (Stam, 2019). Pipe smoking was a common habit that was associated with masculine identity and sociability, together with the consumption of alcohol (Brongers 1964). During excavations of the cemetery at Middenbeemster, eleven 17^th^-19^th^ century clay pipe fragments were recovered from a ditch in the cemetery boundaries (Hakvoort, 2013). There are also multiple contemporaneous sites that have been excavated in the Beemster that have yielded abundant clay pipe fragments (Schabbink, 2020). Furthermore, advertisements indicate that there was also a cigar manufacturing company in the Beemster, making cigars also available to the local population (Inskip et al., n.d.). Snuff and tobacco were also used as ingredients in medical remedies.

Tobacco was economically significant in the post-medieval period and as a result there are plentiful historical sources on its import, processing and use. The kingdoms of Castille and Portugal had the earliest colonies and were exporting tobacco to Europe by the end of the 16^th^ century (Norton, 2008). In the early 17^th^ century, all tobacco imports came through Seville (Goodman, 1993) which also became the leading producer of European snuff. In contrast to the Netherlands, the inhabitants of Iberia were renowned for their cigars and snuff, which likely relates to their early encounters with indigenous peoples who used tobacco in this form (Goodman, 1993). However, depictions of tobacco pipe smoking (Norton, 2008) and the finding of tobacco pipes at Post-Medieval and Modern sites showing that people used it in this form (Cortes Bárcena, 2013; de Heredia Bercero et al., 2012). The graves at the CMB site were found to contain kaolin clay pipes, in addition to coins, rosaries, crosses, medals, remains of ceramics, glass, metal, nails, and fragments of stained glass. Among the many clay pipes recovered, one fragment contained a type of decoration suggesting that it may have originated from a pipe of Dutch manufacture. These clay pipes became popular in the region in the 17th century (Domínguez Ballesteros et al., 2018).

### DNA Extraction

For each Middenbeemster dental calculus sample, 10-30 mg were subsampled for extraction into 1.5 mL tubes. To remove surface contamination, 1 ml of 0.5M EDTA was added to each sample and mixed by vortexing for 20 seconds, followed by 15 minutes of incubation with rotation. The sample solutions were then briefly centrifuged for 1 minute at 6000 rpm (batches 1-4) or 13000 rpm (batch 5) to pellet the calculus fragments, and the supernatant was removed.

To decalcify the decontaminated dental calculus, the resulting pellet was resuspended in 1 ml of 0.5 m EDTA and vortexed for 20 seconds. All samples were then incubated with rotation for 4.75 hours (batch 5), 7 hours (batch 2) or overnight (batches 1, 3, 4). To each sample, 100 μl Proteinase K (30 units/mg) was added, while the controls received 50 μl. All tubes were incubated at 55°C for 5.5 hours (batch 1), 6.5 hours (batch 4), 7.5 hours (batch 3) or overnight (batches 2, 5). There was only enough Qiagen Proteinase K to add to Batch 1-3, therefore Batch 4 and 5 needed to be treated with Invitrogen Proteinase K that has been dissolved in 2.5 ml 99.5% glycerol, 0.5 ml Tns-HCl (100 mM), 0.1 ml CaCl (1M) and 1.9 ml H2O. The samples were held at room temperature and further incubated and digested for five days.

The extraction of the MID DNA was performed using the phenol:chloroform:isoamyl alcohol (25:24:1) method. Solutions B1 and B2 consisted of 375 μl phenol and 375 μl chloroform:isoamylalcohol (phenol:chloroform:isoamlyalcohol 25:24:1), B3 of 750 chloroform: isoamylalcohol. The samples were centrifuged at 13000 rpm for 5 minutes. The supernatant was transferred to solution B1 and the pellet was stored at −20°C. The B1 mix was incubated while being rotated for 1 minute. After a centrifugation step at 13000 rpm for 5 minutes, the aqueous phase was transferred to solution B2, and the rotation and centrifugation step was repeated. The organic phase of B1 was stored at −20°C. Again, the aqueous phase of B2 was transferred to B3, the mixture rotated and centrifuged and the organic phase of B2 stored at −20°C.

The extracted DNA was isolated by silica column-based purification. A MinElute Zymo reservoir with 13 ml PB buffer was placed in a 50 ml falcon tube. The sample was transferred to the PB buffer in the column and centrifuged at 1500 g for 4 minutes, then rotated for 2 minutes. The MinElute column was removed from the reservoir and transferred in a clean collection tube. The column was dry spun at 6000 rpm for 1 minute, and the flow-through was discarded. The DNA containing membranes were washed twice by adding 750 μl PE buffer and centrifugation at 6000 rpm for 1 minute. Each time, the flow-through was discarded. The column underwent another dry spin at 13000 rpm for 1 minute. The MinElute column was transferred to a clean collection tube. A volume of 30 μl EB buffer was added to the center of the filter of the column and incubated for 5 minutes. To elute the DNA from the column, the columns were centrifuged at 13000 rpm for 1 minute. The flow-through was collected, quantified using a Qubit fluorometer and stored at −20°C.

DNA extractions from the CMB, ELR, and IVE sites were performed following the published protocol “Ancient DNA Extraction from Dental Calculus” (Aron et al., 2020a). A single extraction blank, which included water instead of a sample, was included in each extraction batch.

### Library building and Sequencing

Library preparation, indexing, amplification, and pooling for all extraction sets (MID, CMB, ELR, IVE) was identical. Library preparation was performed following the published protocol “Half-UDG treated double-stranded ancient DNA library preparation for Illumina sequencing” (Aron et al., 2020b), including a single library blank per batch, which included water instead of sample extract. Indexing was performed following the published protocol “Illumina double-stranded DNA dual indexing for ancient DNA V.2” (Stahl et al., 2021). Final amplification and pooling were performed following the published protocol “Amplification and Pooling” (Aron and Brandt, 2020).

MID and CMB calculus libraries were pooled in equimolar amounts, and blank libraries pooled in equimolar amounts that were one fifth the concentration of calculus libraries. All libraries, extraction blank libraries, and library build blank libraries were pooled together. Sequencing of the pooled libraries was performed on two flow cells on an Illumina NextSeq500, with 2×75bp chemistry to a depth of ∼8M reads per calculus library and ∼2M reads per blank library (Supplemental Table S1). Sequencing of the IVE and ELR libraries was performed on three independent NextSeq500 runs with 2×150 chemistry for 2 runs, and 2×75bp chemistry for the third (Supplemental Table S1). Fastq files for independent sequencing runs were merged per sample in the data processing steps described below.

### Data processing

All raw data were processed using the nf-core/eager pipeline (Fellows Yates et al., 2021a), version 2.1.0. This included quality checks with FastQC (Andrews, 2010), adapter trimming, read collapsing, and quality filtering with AdapterRemoval (Schubert et al., 2016), and mapping against the human genome (HG19) with bwa aln -n 0.02 -l 1024 (Li and Durbin, 2009) and samtools (Danecek et al., 2021) to remove human reads. The human-mapped reads were not used for any analyses. Taxonomic profiling was performed with MALT v. 0.4.0 (Herbig et al., 2016; Vågene et al., 2018). The command to run this can be found in the github repository https://github.com/ivelsko/smoking_calculus/02-scripts.backup/smokers_calc_notes.txt. Metadata for data processing for data produced for this study are in Supplemental Table S6.

### Processing of comparative data

These published historic dental calculus samples were downloaded from ENA and processed with the nf-core/eager pipeline (Fellows Yates et al., 2021a), described above. For analyses of pre- and post-tobacco introduction populations, post-medieval individuals from RAD, MID, and CMB were treated as smokers and medieval individuals from KIL, ELR, and IVE were treated as non-smokers, since no tobacco products were available in Europe during the lifetime of these individuals. Metadata for data processing for published data used in this study are in Supplemental Table S7.

The published species table from Supplemental Table S1 in (Fagernäs et al., 2022), was used as input for assessing the distribution of *Streptococcus* species across the dentition of four individuals. This table was generated by processing the data in a similar manner to the rest of this study. In brief, the data was processed with the nf-core/eager pipeline with identical settings as used here, with the exception that a different version of the human genome was used as a reference for mapping. All reads that did not map to the human genome were profiled with MALT v 0.4.0 (Herbig et al., 2016; Vågene et al., 2018) within the nf-core/eager pipeline, using the same settings and the same RefSeq-based database as used in this study, and the species table was produced in MEGAN6 CE from the MALT output.

### Taxonomic profiling and decontamination

All remaining reads that did not map to the human genome were taxonomically profiled with two profilers: MetaPhlAn3 (Beghini et al., 2021) run as a stand-alone program, and MALT v. 0.4.0 (Herbig et al., 2016; Vågene et al., 2018) within the nf-core/eager pipeline. All microbial species diversity and correlation analyses, as well as cuperdec (Fellows Yates et al., 2021b) preservation analysis, were performed with the MetaPhlAn3 species table. The commands to generate the table can be found in /mnt/archgen/microbiome_calculus/smoking_calculus/02-scripts.backup/009-metaphlan3_mid.Snakefile. The MALT species table was used strictly for SourceTracker analysis and for investigating *Streptococcus* species distributions, as we found a taxonomic gradient across samples resulting in an artifact during diversity analysis (described below in the section “*Taxonomic gradient investigation*”). MALT was run within the nf-core/eager pipeline with default settings. The database used was the custom RefSeq database described in Fellows Yates (Fellows Yates et al., 2021b). The output rma6 files from MALT were imported to MEGAN6 CE v. 6.18.0 (Huson et al., 2016) with the “comparison” mode, and a species-level table with read counts was exported as a tsv file for all downstream analyses. The MetaPhlAn3 species table is presented as Supplemental Table S8, and the MALT species table can be found on the github repository as an RData file https://github.com/ivelsko/smoking_calculus/05-results.backup/Taxonomy_tables.RData or as a tsv file https://github.com/ivelsko/smoking_calculus/05-results.backup/malt_refseq_species.tsv

Poorly preserved dental calculus samples were identified using the R package cuperdec (Fellows Yates et al., 2021b) using the MetaPhlAn3 table, and were removed from the table for all downstream processing. Potential contaminant taxa were removed from the taxonomic table using the R package decontam (Davis et al., 2018), with extraction blanks and library blanks from this study, as well as femur samples from (Fellows Yates et al., 2021b), as controls. Scripts for these steps can be found in the github repository: https://github.com/ivelsko/smoking_calculus/02-scripts.backup/ in the files MID_mpa3_cuperdec.Rmd, MID_mpa3_decontam.Rmd.

SourceTracker (Knights et al., 2011) was used to determine the proportion of species in each sample that come from a set of authentic oral and potential contamination sources. SourceTracker was run using the MALT species table as input because species profiles of the source sample generated with MALT using the RefSeq database from (Fellows Yates et al., 2021b) were available, negating the need to download, process, and taxonomically profile the source raw data. Species tables for all sources except modern calculus were obtained from the table Evolution-Comparison_MEGAN_20190410-ex_absolute_species_prokaryotes_summarised_refseq.txt from the github page of Fellows Yates, et al. (Fellows Yates et al., 2021b), and the following sources were used: modern dental calculus, supragingival plaque, subgingival plaque, rural gut, urban gut, skin, archaeological bone, and sediment. Modern calculus source data was obtained from Velsko, et al. (Velsko et al., 2019) and Fellows Yates, et al. (Fellows Yates et al., 2021b) and was processed through nf-core/eager and MALT with the ancient calculus for this study. Scripts for these steps can be found in the github repository: https://github.com/ivelsko/smoking_calculus/02-scripts.backup/ in the files 001-shotgun_sourcetracker_high_rare.sh and MID_tax_sourcetracker.Rmd.

### Diversity analyses and metadata comparisons

The MetaPhlAn3 species table was used for all diversity analyses. The Shannon index was calculated in R using the diversity function in the package vegan (OKSANEN and J, 2007). Kruskal-wallace tests were performed using the R package rstatix (Kassambara, 2020). A PCA was calculated in R on a CLR-transformed species table using the package mixOmics (Rohart et al., 2017). PERMANOVA was run on the PCA using the function adonis2 in the R package vegan (OKSANEN and J, 2007). Batch effects within the MID sample set were investigated by coloring the points in a PCA plot by extraction batch, but no clustering based on batch was observed (Supplemental Figure S4). https://github.com/ivelsko/smoking_calculus/06-publication/main_figures/ in MID_mpa3_alphadiv.Rmd and MID_mpa3_betadiv.Rmd.

### Subsampled datasets

The effects of library sequencing depth and average read length on the number of species detected were investigated by down-sampling the full libraries in two ways. For the first set, we randomly subsampled all libraries with > 10M reads down to 10M reads using seqtk and setting the seed to -s10000, while leaving all libraries with < 10M reads untouched (Sub 10M set). For the second set, we subsampled all libraries to include only reads ≤ 75bp in length (Sub 75bp set) using bioawk -c fastx ‘{if (length($seq) < 76){print “@“$name” “$comment”\n“$seq”\n+\n“$qual}}’ <library>. The full libraries that were downsampled were those that had been processed by nf-core/eager (adapter-trimmed and quality-filtered, collapsed, mapped against the human genome, and had human reads removed), and were profiled by MetaPhlAn3 for full analysis. Both subsetted datasets were profiled with MetaPhlAn3 as described above for the full set. The total number of reads, the average read length, and the average GC content of the libraries for both subsetted datasets were calculated from FASTQC (Andrews, 2010) using multiqc (Ewels et al., 2016) (Supplemental Tables S6, S7) and compared with the full set. Alpha-diversity metrics Observed species (number of species) and Shannon index were calculated as described for the full set in Diversity analyses and metadata comparisons (Supplemental Table S9). Scripts for these steps can be found in the github repository: https://github.com/ivelsko/smoking_calculus/02-scripts.backup/ in the files MID_mpa3_alphadiv_sub10M.Rmd, MID_mpa3_alphadiv_sub75bp.Rmd, MID_mpa3_sp_counts_sub10M_sub75bp.Rmd, and MID_sub10M_sub75bp_numbers.Rmd.

### Canonical correlation analysis

Canonical correlation analysis was performed to look for correlations between different metadata categories and between metadata and principal component loadings from PCA, as in (Briscoe et al., 2022). Input tables contained selected metadata categories and the PC1 and PC2 loadings from PCA. Canonical correlations were calculated with the function canCorPairs from the R package variancePartition (Hoffman and Roussos, 2021; Hoffman and Schadt, 2016). Statistical tests were performed with the cor.mtest function in the R package corrplot (Wei et al., 2013), and correlation matrix plots were generated with the function corrplot in the same package. To focus on the strongest correlations, we considered only correlations ≥ 0.4 with a significance of p ≤ 0.05. Scripts for these steps can be found in the github repository: https://github.com/ivelsko/smoking_calculus/02-scripts.backup/MID_mpa3_PC_metadata_corr.Rmd and https://github.com/ivelsko/smoking_calculus/06-publication/main_figures/ in Figure_2/Figure_2.Rmd and Figure_5/Figure_5.Rmd.

### Taxonomic gradient investigation

Beta diversity analysis PCA plots based on the MALT RefSeq table showed a distinctive horseshoe shape, indicating a taxonomic gradient (Morton et al., 2017). The taxa responsible for this were investigated by examining the species with strongest loadings in PC1 positive space and PC1 negative space. When the taxon abundance was plotted as a heatmap with the samples ordered by loading in PC1, a diagonal gradient was observed (Supplemental Figure S5). Samples at one end of the PCA horseshoe were characterized by early colonizer taxa, while those at the other end were characterized by poorly characterized taxa, mostly isolated from non-human sources such as environment and anaerobic sludge digestors. Since filtering could not remove these environmental taxa, which are likely mis-assignments, we used the MetaPhlan3 input table instead for all downstream diversity analyses. MetaPhlan2 was shown to produce a highly accurate species profile on ancient dental calculus data, with very low numbers of false positive taxa identified (Velsko et al., 2018). Scripts for these steps can be found in the github repository: https://github.com/ivelsko/smoking_calculus/02-scripts.backup/MID_tax_horseshoe.Rmd.

### HUMAnN3 functional analysis

Potential metabolic functional profiles were generated with HUMAnN3 (Beghini et al., 2021) using default parameters. We used the pathway abundance table, which was converted from reads per kilobase to copies per million with the humann3 helper script humann_renorm_table.py. The total pathway assignment per sample was used in analysis, and not the species-specific assignments per pathway. A PCA was performed on the pathway abundance table using MixOmics as described for the species tables above. A canonical correlation analysis was performed on the principal component loadings, the metadata, and the loadings from the species PCA, as described above in Canonical Correlation Analysis. Scripts for these steps can be found in the github repository: https://github.com/ivelsko/smoking_calculus/02-scripts.backup/MID_humann3_paths.Rmd.

### Streptococcus species distributions

Dental calculus profiles are distinct from dental plaque, but can be highly variable between individuals. An earlier study (Fellows Yates 2021) noted a distinct clustering of human calculus samples with a *Streptococcus* species profile resembling that found in chimpanzees, yet was not able to associate this profile with any metadata due to low sample numbers. We used our large cohort from a single population in a single cemetery to investigate if this trend is observable in a dataset where the reasons for this might be investigated. Using the species table from MALT RefSeq profiling, we assigned each *Streptococcus* species to a group as used in (Fellows Yates et al., 2021b): Sanguinis, Mitis, Salivarius, Anginosus, Bovis, Pyogenic, Mutans, unknown, NA, which come from (Richards et al., 2014).

To be able to group *Streptococcus* genomes that had reads aligned by MALT but were not assigned a species name, we ran ‘dRep cluster’ on all *Streptococcus* genomes with hits in any sample. *Streptococcus* isolates with no species designation were then assigned to a group based on which clade defined by Richards, et al. 2014, they fell into in the dRep primary clustering phylogeny. We calculated the proportion of reads from *Streptococcus* species in each of these groups out of all *Streptococcus* species-assigned reads in each sample. Further, within each sample, we calculated the proportion of reads that were assigned to any species in the genus *Streptococcus* vs. all other genus assignments. Scripts for these steps can be found in the github repository: https://github.com/ivelsko/smoking_calculus/02-scripts.backup/ in the files 010-dRep_Strep_nunknowns.Snakefile and MID_Strep_groups.Rmd and DA_Strep_groupt.Rmd.

### Additional plotting aspects

Plots were arranged in grids using cowplot (Wilke, 2020) or patchwork v1.1.0 (Pedersen, 2017) in R. Metadata plots used ggpointgrid (Schmid, 2022). Significance on plots was indicated with the R package ggsignif (Ahlmann-Eltze and Patil, 2021). A map of Europe with the sites of MID, CMB, and comparative data sites was generated in R using the packagtes sf (Pebesma, 2018), rnaturalearth (South, 2017a), rnaturalearthdata (South, 2017b)(), rgeos (Bivand et al., 2017)(), and maps (Becker et al., 2021). Scripts to generate all main and supplemental figures can be found in their respective folders in the github repository: https://github.com/ivelsko/smoking_calculus/06-publication/.

## Data Availability

All data generated for this study has been uploaded to the European Nucleotide Archive under accession **PRJEB52394**. Scripts for analysis can be found on the github repository https://github.com/ivelsko/smoking_calculus.

## Author Contributions

I.M.V., S.I., M.I.G.C., M.H., and C.W. designed the study. S.I. and M.I.G.C. performed the osteological analyses. L.S. and K.Z. performed the laboratory analyses. I.M.V. and S.I. performed data analysis. M.H., M.I.G.C., M.R.S., L.B.L.E, J.M.M.G, D.G.V., A.C.P.R., D.C.S.G. and K.Z. provided materials and resources including performing calculus sampling. I.M.V., S.I., and C.W. wrote the manuscript, with contributions from all coauthors.

## Acknowledgments

We thank Antje Wissgot for assistance with sequencing. We thank the Historisch Genootschap Beemster for access to the Middenbeemster skeletal collection. We also like to thank the Middenbeemster community for allowing access to the remains of their ancestors. We thank Qark Arqueología (Eder Domínguez-Ballesteros, Leandro Sánchez) for access to the calculus samples from Convento de los Mercedarios de Burtzeña, Spain. We thank Julio Trelis/Ayuntament de Crevillent for access to the calculus samples from El Raval, Spain. This research was supported by the Werner Siemens Stiftung (I.M.V. and C.W.), the Deutsche Forschungsgemeinschaft (DFG, German Research Foundation) under Germany’s Excellence Strategy EXC 2051 Project-ID 390713860 (I.M.V. and C.W.), the Max Planck Society, and the UKRI AHRC FLF grant MR/T022302/1 (S.I.).), and the Government of the Basque Country grant POS_2020_1_0006 (M.I.G.C.). DCSG acknowledges funding from the Generalitat Valenciana (CIDEGENT 2019/061) and the Spanish government (EUR2020-112213). DCSG acknowledges funding from the Generalitat Valenciana (CIDEGENT 2019/061) and the Spanish government (EUR2020-112213).

## Supporting Information

### List of Supplemental Tables in Excel sheets

**Table S1.** Oral pathology metadata for samples sequenced for this study. (Excel sheet)

**Table S5.** Oral pathology metadata for published samples included in this study. (Excel sheet)

**Table S6.** Extraction and library metadata for samples sequenced for this study (Excel sheet)

**Table S7.** Extraction and library metadata for published samples included in this study (Excel sheet)

**Table S8.** MetaPhlAn3 taxonomy table for all samples included in this study. (Excel sheet)

**Table S2.**
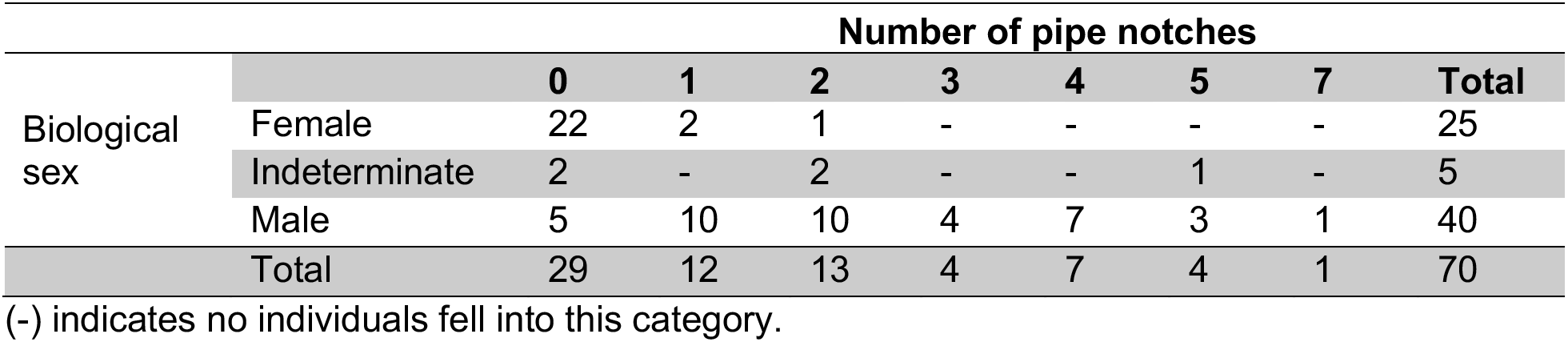
Number of pipe notches in individuals from Middenbeemster.

|  |  | Number of pipe notches |  |  |  |  |  |  | Total |
| --- | --- | --- | --- | --- | --- | --- | --- | --- | --- |
|  |  | 0 | 1 | 2 | 3 | 4 | 5 | 7 |  |
| Biological sex | Female | 22 | 2 | 1 | - | - | - | - | 25 |
|  | Indeterminate | 2 | - | 2 | - | - | 1 | - | 5 |
|  | Male | 5 | 10 | 10 | 4 | 7 | 3 | 1 | 40 |
|  | Total | 29 | 12 | 13 | 4 | 7 | 4 | 1 | 70 |

(-) indicates no individuals fell into this category.

**Table S3.**
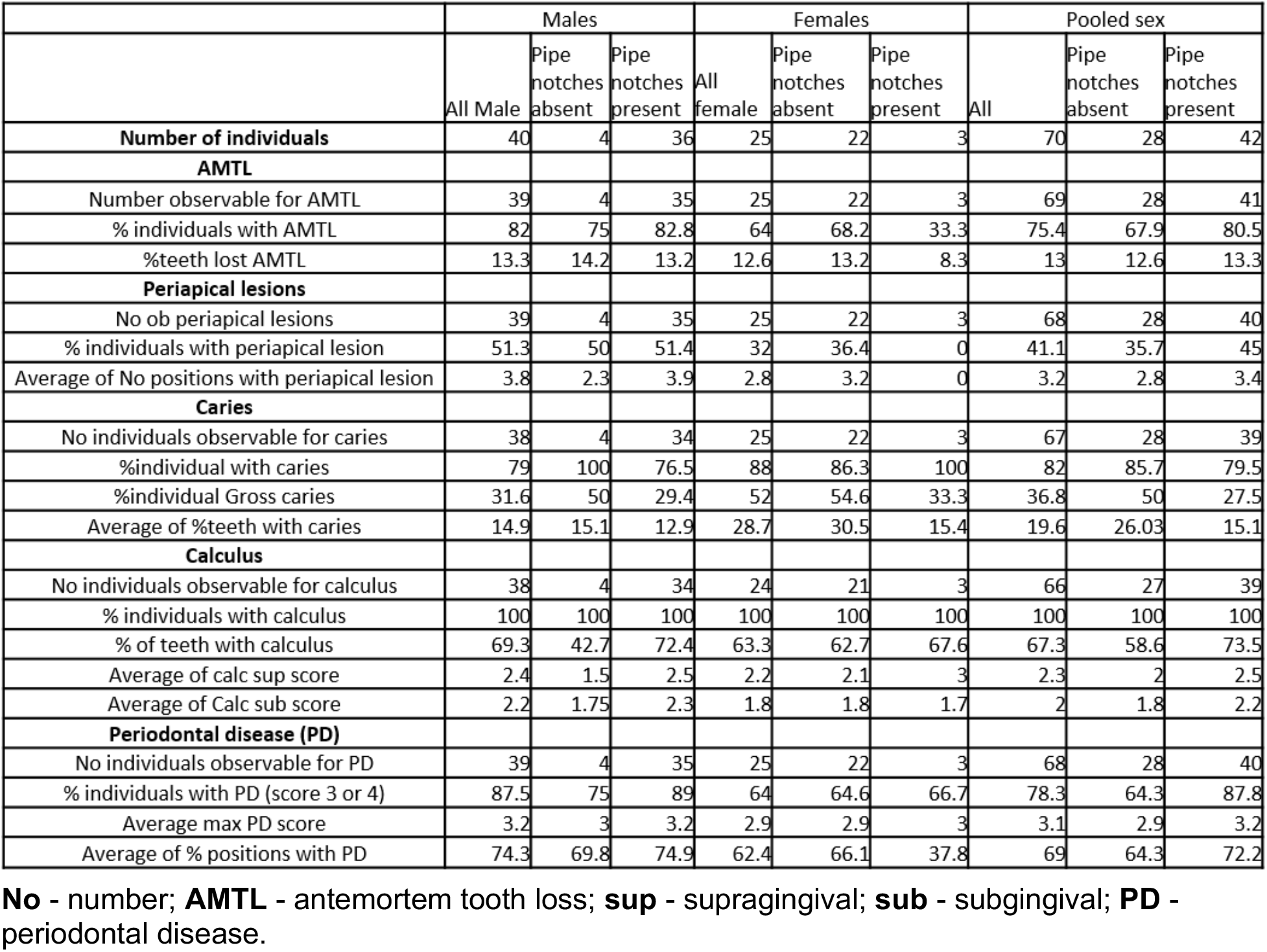
Numbers of observations and prevalence rates for oral pathologies for males, females, and a pooled sex group.

|  | Males |  |  | Females |  |  | Pooled sex |  |  |
| --- | --- | --- | --- | --- | --- | --- | --- | --- | --- |
|  | All Male | Pipe notches absent | Pipe notches present | All female | Pipe notches absent | Pipe notches present | All | Pipe notches absent | Pipe notches present |
| **Number of individuals** | 40 | 4 | 36 | 25 | 22 | 3 | 70 | 28 | 42 |
| **AMTL** |  |  |  |  |  |  |  |  |  |
| Number observable for AMTL | 39 | 4 | 35 | 25 | 22 | 3 | 69 | 28 | 41 |
| % individuals with AMTL | 82 | 75 | 82.8 | 64 | 68.2 | 33.3 | 75.4 | 67.9 | 80.5 |
| %teeth lost AMTL | 13.3 | 14.2 | 13.2 | 12.6 | 13.2 | 8.3 | 13 | 12.6 | 13.3 |
| **Periapical lesions** |  |  |  |  |  |  |  |  |  |
| No ob periapical lesions | 39 | 4 | 35 | 25 | 22 | 3 | 68 | 28 | 40 |
| % individuals with periapical lesion | 51.3 | 50 | 51.4 | 32 | 36.4 | 0 | 41.1 | 35.7 | 45 |
| Average of No positions with periapical lesion | 3.8 | 2.3 | 3.9 | 2.8 | 3.2 | 0 | 3.2 | 2.8 | 3.4 |
| **Caries** |  |  |  |  |  |  |  |  |  |
| No individuals observable for caries | 38 | 4 | 34 | 25 | 22 | 3 | 67 | 28 | 39 |
| %individual with caries | 79 | 100 | 76.5 | 88 | 86.3 | 100 | 82 | 85.7 | 79.5 |
| %individual Gross caries | 31.6 | 50 | 29.4 | 52 | 54.6 | 33.3 | 36.8 | 50 | 27.5 |
| Average of %teeth with caries | 14.9 | 15.1 | 12.9 | 28.7 | 30.5 | 15.4 | 19.6 | 26.03 | 15.1 |
| **Calculus** |  |  |  |  |  |  |  |  |  |
| No individuals observable for calculus | 38 | 4 | 34 | 24 | 21 | 3 | 66 | 27 | 39 |
| % individuals with calculus | 100 | 100 | 100 | 100 | 100 | 100 | 100 | 100 | 100 |
| % of teeth with calculus | 69.3 | 42.7 | 72.4 | 63.3 | 62.7 | 67.6 | 67.3 | 58.6 | 73.5 |
| Average of calc sup score | 2.4 | 1.5 | 2.5 | 2.2 | 2.1 | 3 | 2.3 | 2 | 2.5 |
| Average of Calc sub score | 2.2 | 1.75 | 2.3 | 1.8 | 1.8 | 1.7 | 2 | 1.8 | 2.2 |
| **Periodontal disease (PD)** |  |  |  |  |  |  |  |  |  |
| No individuals observable for PD | 39 | 4 | 35 | 25 | 22 | 3 | 68 | 28 | 40 |
| % individuals with PD (score 3 or 4) | 87.5 | 75 | 89 | 64 | 64.6 | 66.7 | 78.3 | 64.3 | 87.8 |
| Average max PD score | 3.2 | 3 | 3.2 | 2.9 | 2.9 | 3 | 3.1 | 2.9 | 3.2 |
| Average of % positions with PD | 74.3 | 69.8 | 74.9 | 62.4 | 66.1 | 37.8 | 69 | 64.3 | 72.2 |

**No** - number; **AMTL** - antemortem tooth loss; **sup** - supragingival; **sub** - subgingival; **PD** - periodontal disease.

**Figure S1.**
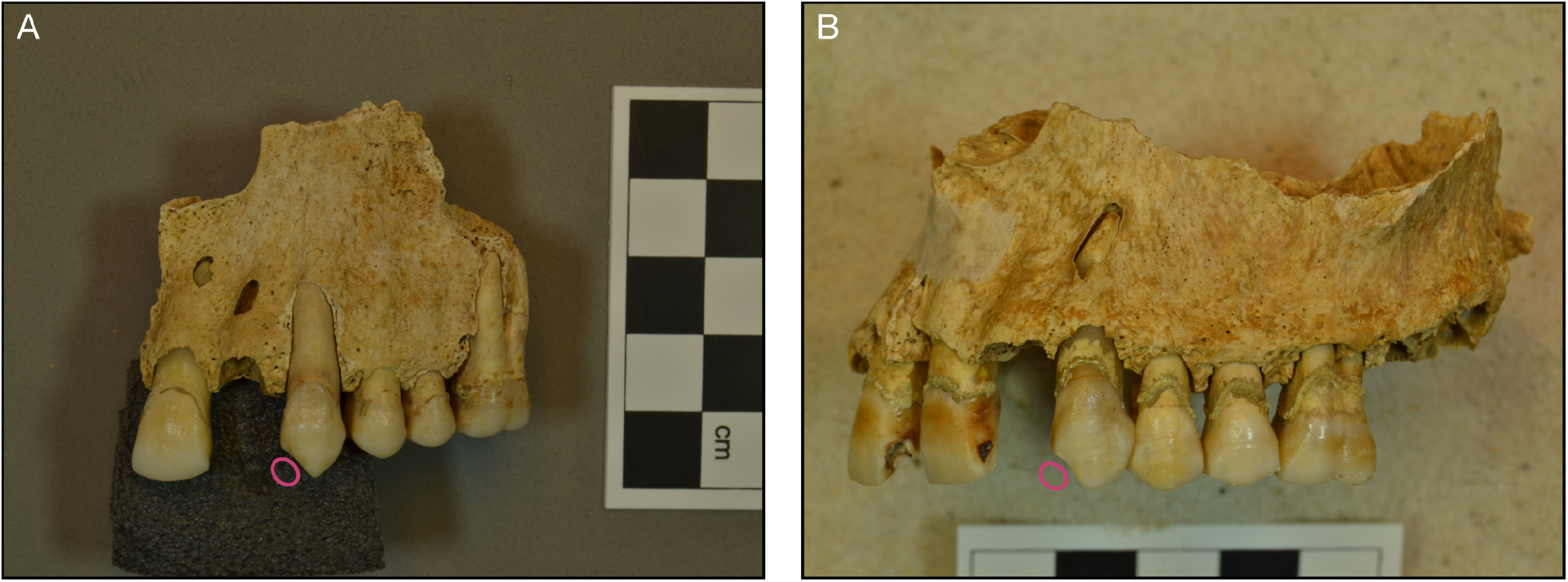
Pipe notches in anterior maxillary dentition of 2 individuals from Convento de los Mercedarios de Burtzeña (CMB). **A.** Individual CMB001. **B.** Individual CMB003. Notch locations are indicated by hollow pink circles. Photo credit: Maite I. Garcia-Collado.

**Figure S2.**
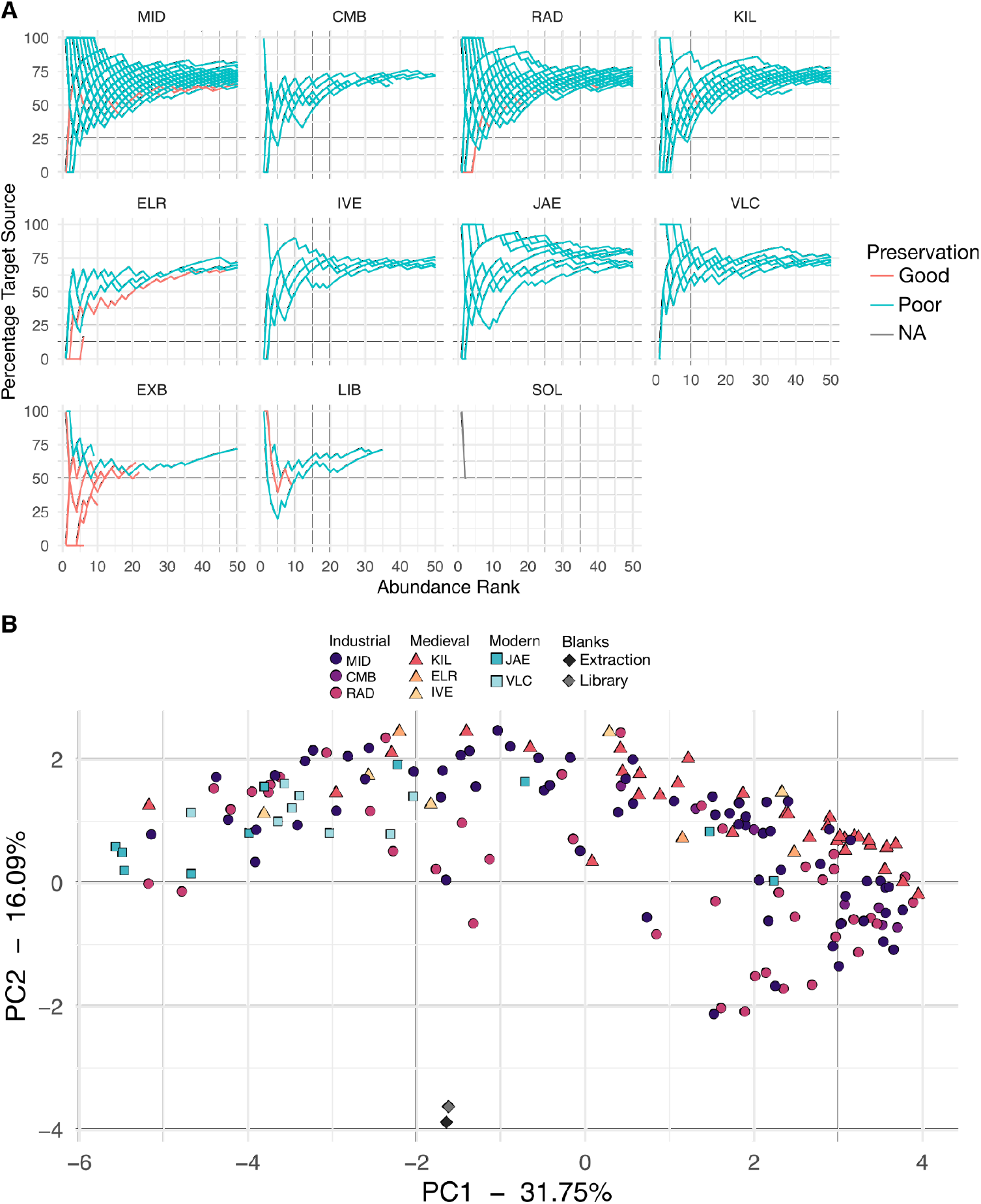
Preservation assessment. A. Cumulative percent decay curves (cuperdec). Curves demonstrating the proportion of species that come from an oral source, ordered by abundance. All samples with red lines were removed from all analyses. B. PCA of samples passing cuperdec preservation cut-offs, as well as extraction and library blanks. Site codes: **MID** - Middenbeemster, **CMB** - Convento de los Mercedarios de Burtzeña, **RAD** - Radcliffe, KIL - Kilteasheen, **ELR** - El Raval, **IVE** - Iglesia de la Virgen de la Estrella, **JAE** - Jaen, **VLC** - Valencia, **EXB** - Extraction blank, **LIB** - library blank, **SOL** - soil.

**Figure S3.**
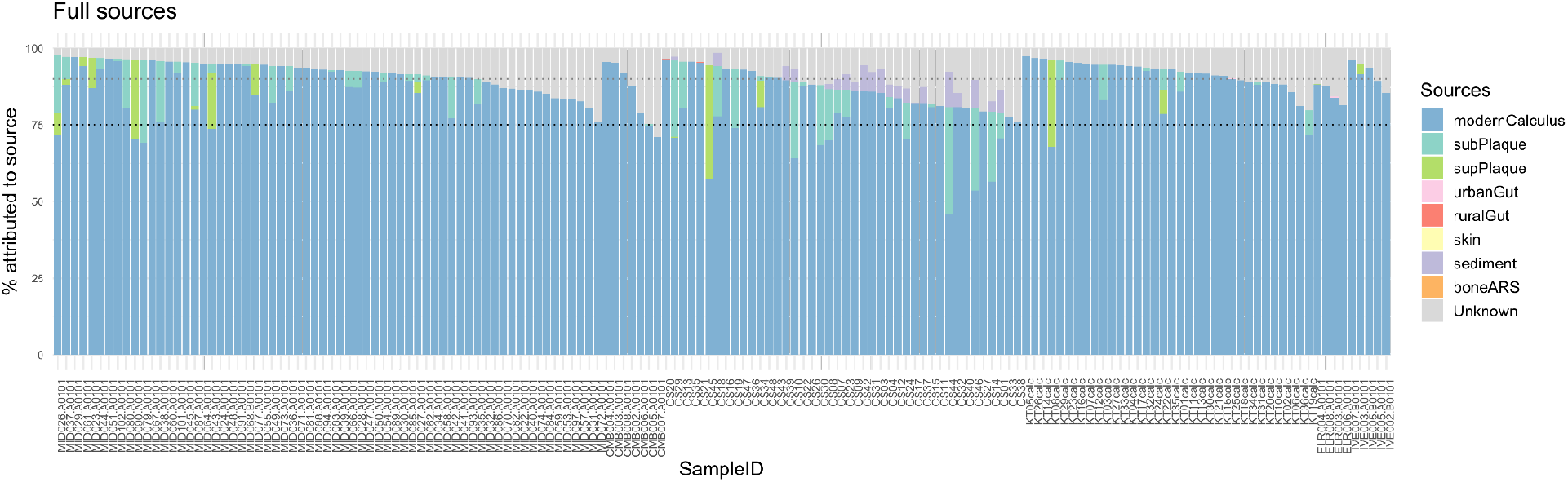
SourceTracker plot showing in each sample the proportion of species assigned to each source. Only a single sample has a proportion below 75% assigned to an oral source (modern calculus, subgingival plaque, supragingival plaque). The input table was from a MALT run that used the RefSeq database from Fellows Yates, et al. 2021. SubPlaque - subgingival plaque, supPlaque - supragingival plaque, boneARS - bones from site Arbulag sum, Mongolia (site code ARS). The modern calculus used as a source is the JAE samples used in this study as comparative samples. The dotted black line indicates 75%, and the dotted gray line indicates 90%.

**Figure S4.**
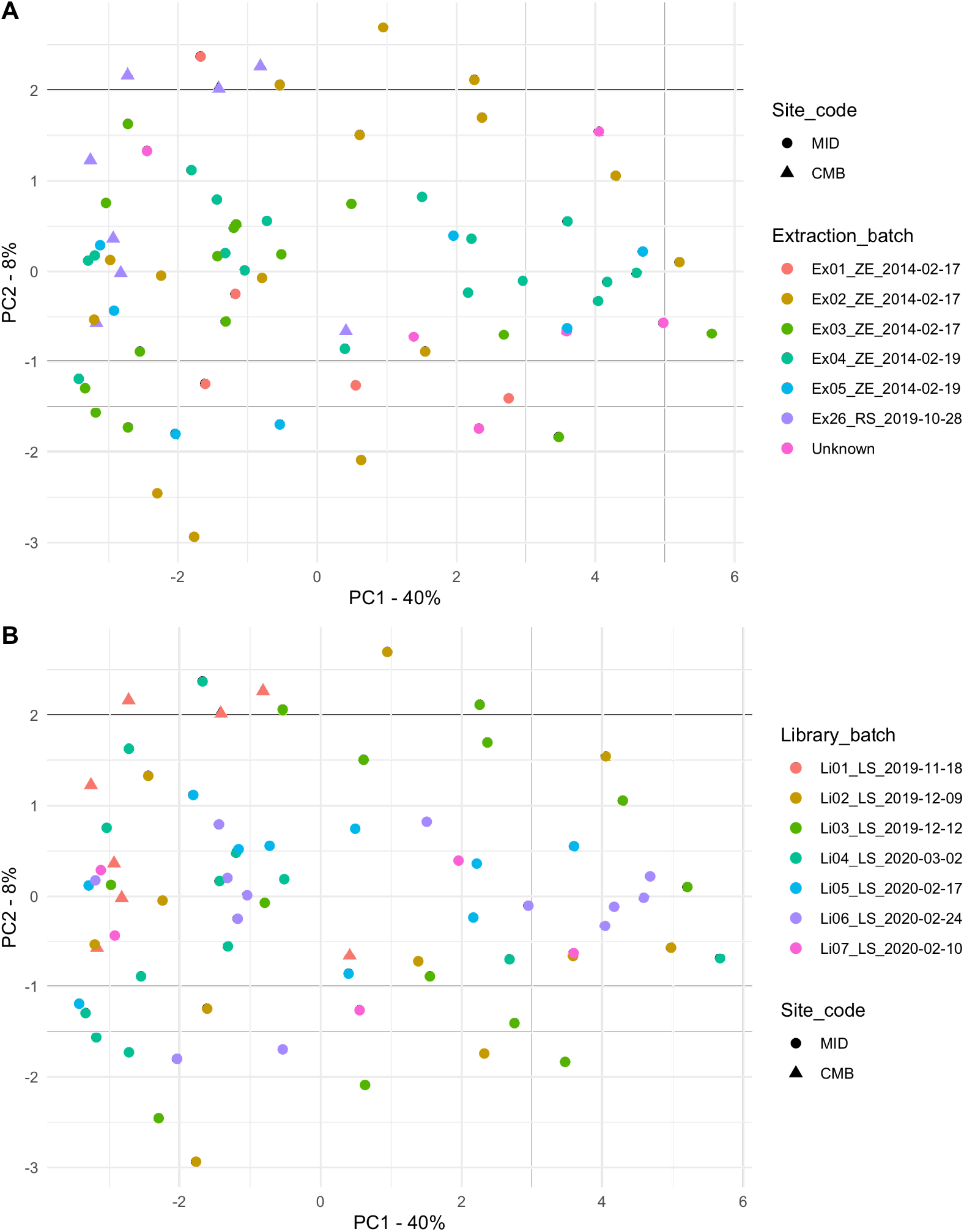
PCA plot of species diversity for Middenbeemster (MID) and Convento de los Mercedarios de Burtzeña (CMB) colored by **A.** extraction batch and **B.** library batch. Samples do not cluster by either extraction or library batch.

**Figure S5.**
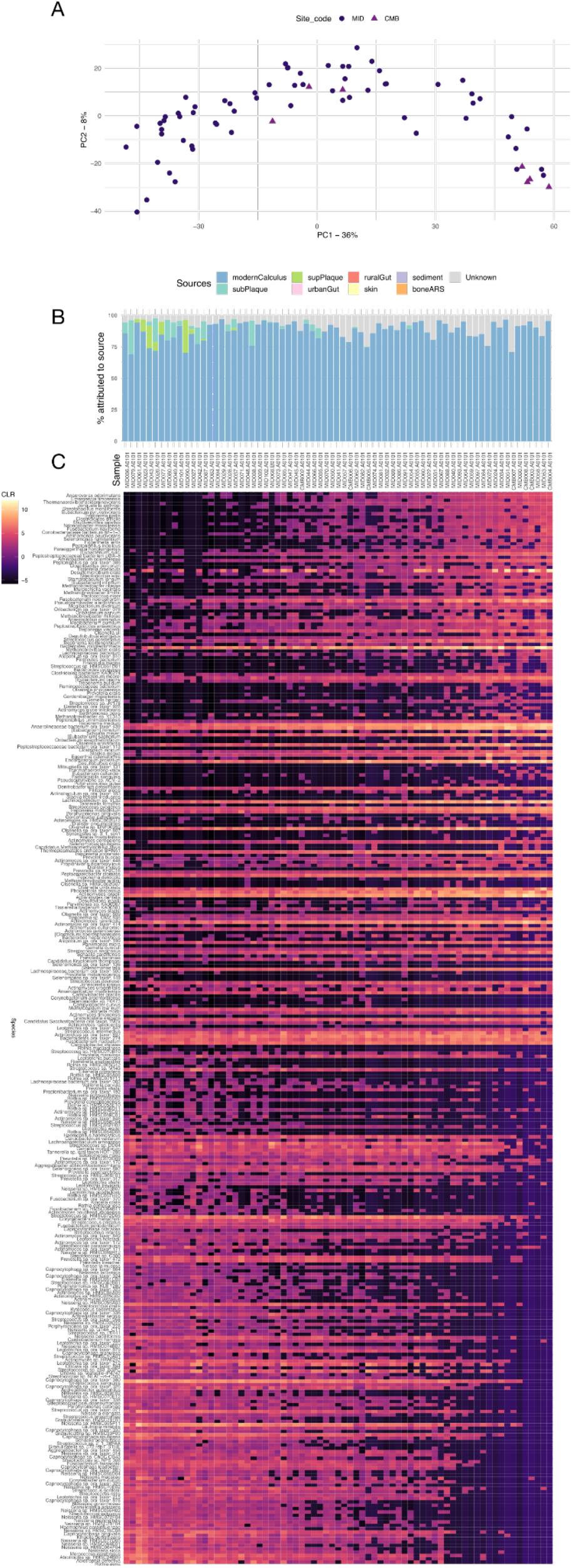
Species gradient in Middenbeemster (MID) and Convento de los Mercedarios de Burtzeña (CMB) calculus samples profiled with MALT. The sample order of plots B and C follows the order of points along PC1 in the PCA shown in panel A. **A.** PCA of read-based species counts result in a horseshoe pattern PCA plot. **B.** SourceTracker results indicate that the samples with the most negative PC1 loadings have higher proportions of species found in supra- and subgingival dental plaque than the other samples. **C.** Heat map showing the CLR-transformed abundance of species present at > 0.01% abundance. A gradient of taxa in samples from one end of the PCA to the other end can be traced from the upper right corner to the lower left corner. Samples with the most negative PC1 loadings have higher proportions of early-colonizer, aerobic and facultative taxa, and lower proportions of late-colonizer, anaerobic taxa than samples with the most positive PC1 loadings, consistent with a higher source contribution of plaque seen in panel B.

**Figure S6.**
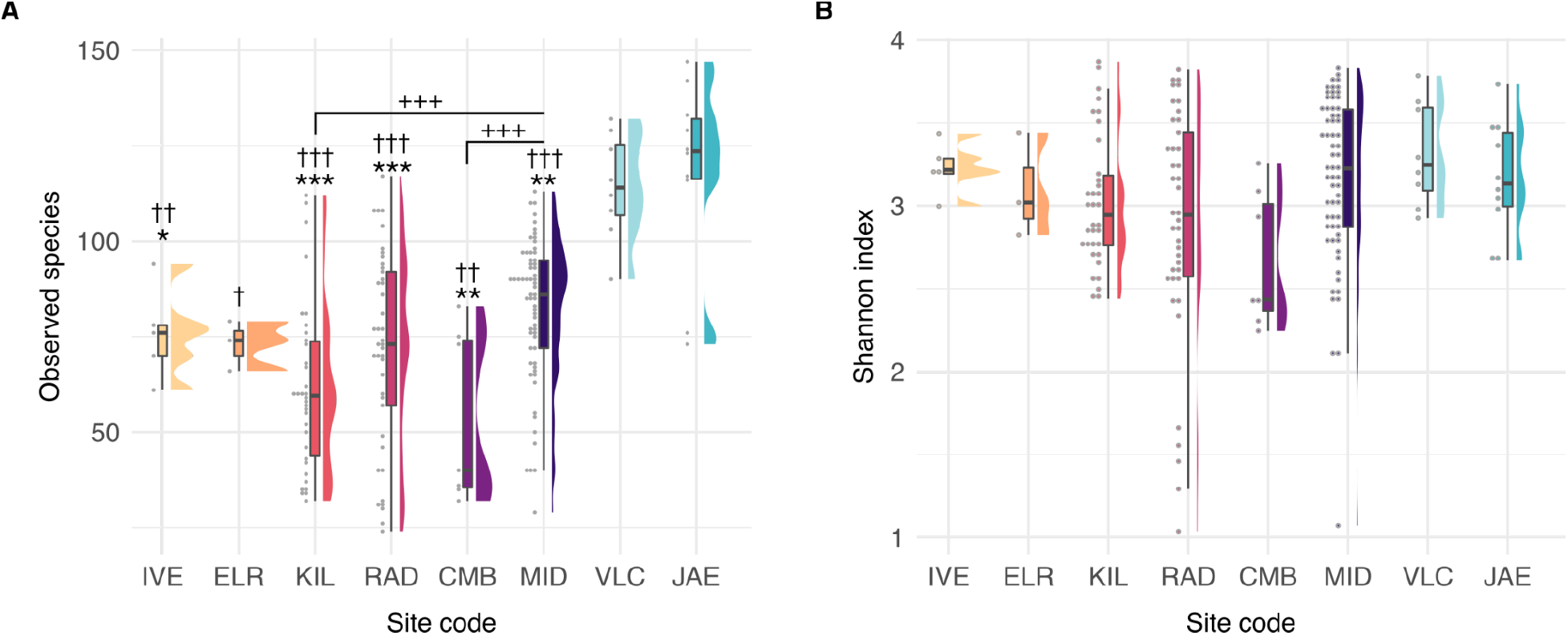
Alpha-diversity of samples grouped by site. A. Number of species. B. Shannon index. *** p < 0.001, ** p < 0.01, * p < 0.05 compared to JAE. ††† p < 0.001, †† p < 0.01, † p < 0.05 compared to VLC. ++ p < 0.01, +++ p < 0.001 compared to MID. No significant differences in Shannon index were detected between sites. Site codes: **IVE** - Iglesia de la Virgen de la Estrella; **ELR** - El Raval; **KIL** - Kilteasheen; **RAD** - Radcliffe; **CMB** - Convento de los Mercedarios de Burtzeña; **MID** - Middenbeemster; **VLC** - Valencia; **JAE** - Jaen.

**Figure S7.**
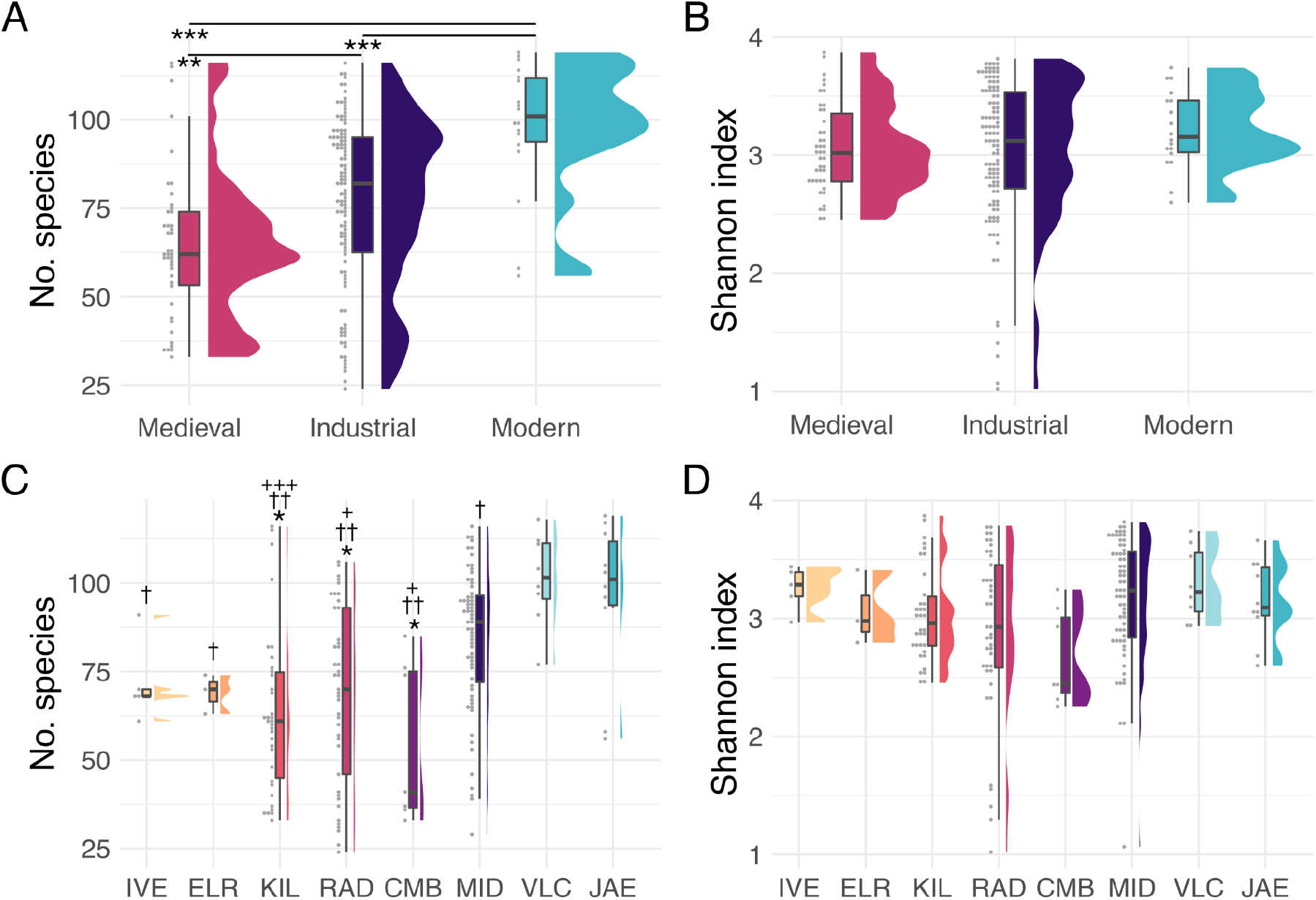
Alpha-diversity after sub-sampling all libraries to a maximum of 10M reads. **A.** Number of species grouped by time period. **B.** Shannon index grouped by time period. **C.** Number of species grouped by site. **D.** Shannon index grouped by site. *** p < 0.001, ** p < 0.01, * p < 0.05 compared to JAE. ††† p < 0.001, †† p < 0.01, † p < 0.05 compared to VLC. + p < 0.05, ++ p < 0.01, +++ p < 0.001 compared to MID. No significant differences in Shannon index were detected between time periods or sites. Site codes: **IVE** - Iglesia de la Virgen de la Estrella; **ELR** - El Raval; **KIL** - Kilteasheen; **RAD** - Radcliffe; **CMB** - Convento de los Mercedarios de Burtzeña; **MID** - Middenbeemster; **VLC** - Valencia; **JAE** - Jaen.

**Figure S8.**
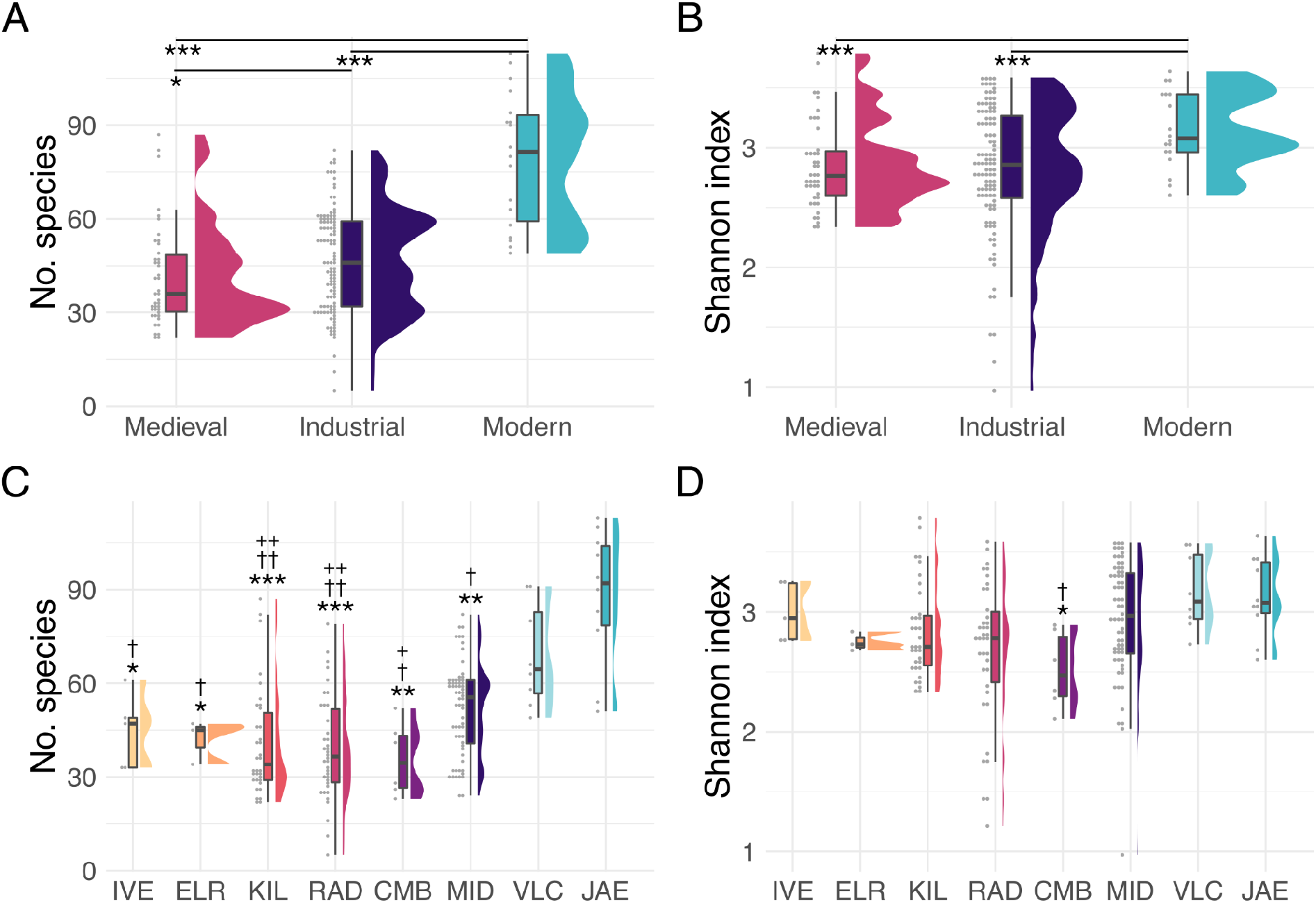
Alpha-diversity after sub-sampling all libraries for only reads 75 bp. **A.** Number of species grouped by time period. **B.** Shannon index grouped by time period. **C.** Number of species grouped by site. **D.** Shannon index grouped by site. *** p < 0.001, ** p < 0.01, * p < 0.05 compared to JAE. ††† p < 0.001, †† p < 0.01, † p < 0.05 compared to VLC. + p < 0.05, ++ p < 0.01, +++ p < 0.001 compared to MID. No significant differences in Shannon index were detected between time periods. Site codes: **IVE** - Iglesia de la Virgen de la Estrella; **ELR** - El Raval; **KIL** - Kilteasheen; **RAD** - Radcliffe; **CMB** - Convento de los Mercedarios de Burtzeña; **MID** - Middenbeemster; **VLC** - Valencia; **JAE** - Jaen.

**Figure S9.**
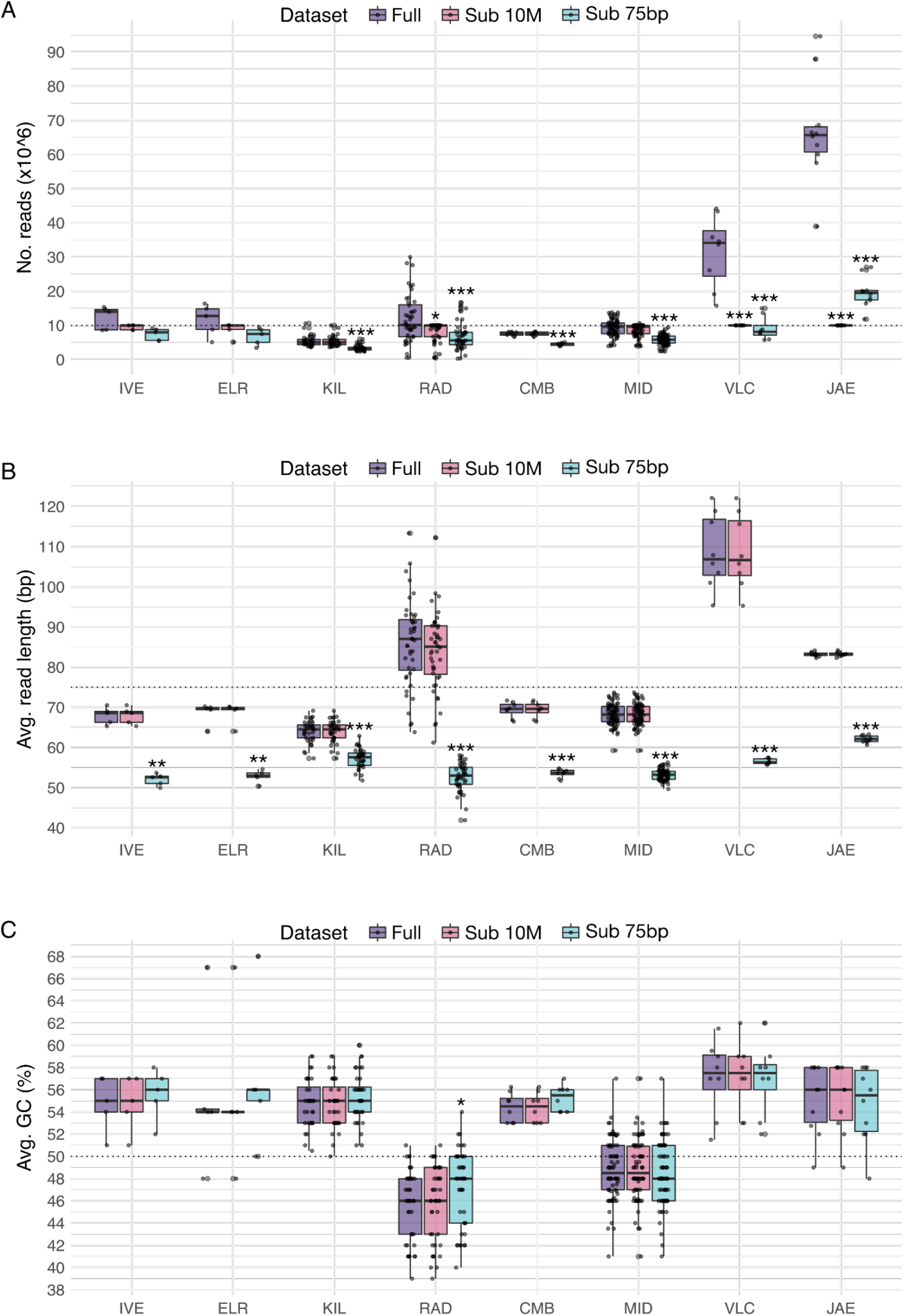
Read characteristics for full and sub-sampled libraries. A. Total number of reads. B. Average read length (bp). C. Average GC content (%). Sub 10M - libraries subsampled to include no more than 10 million reads. Sub 75bp - libraries subsampled to include only reads ≤ 75bp in length. *** p < 0.001, ** p < 0.01, * p < 0.05 compared to the full dataset

**Figure S10.**
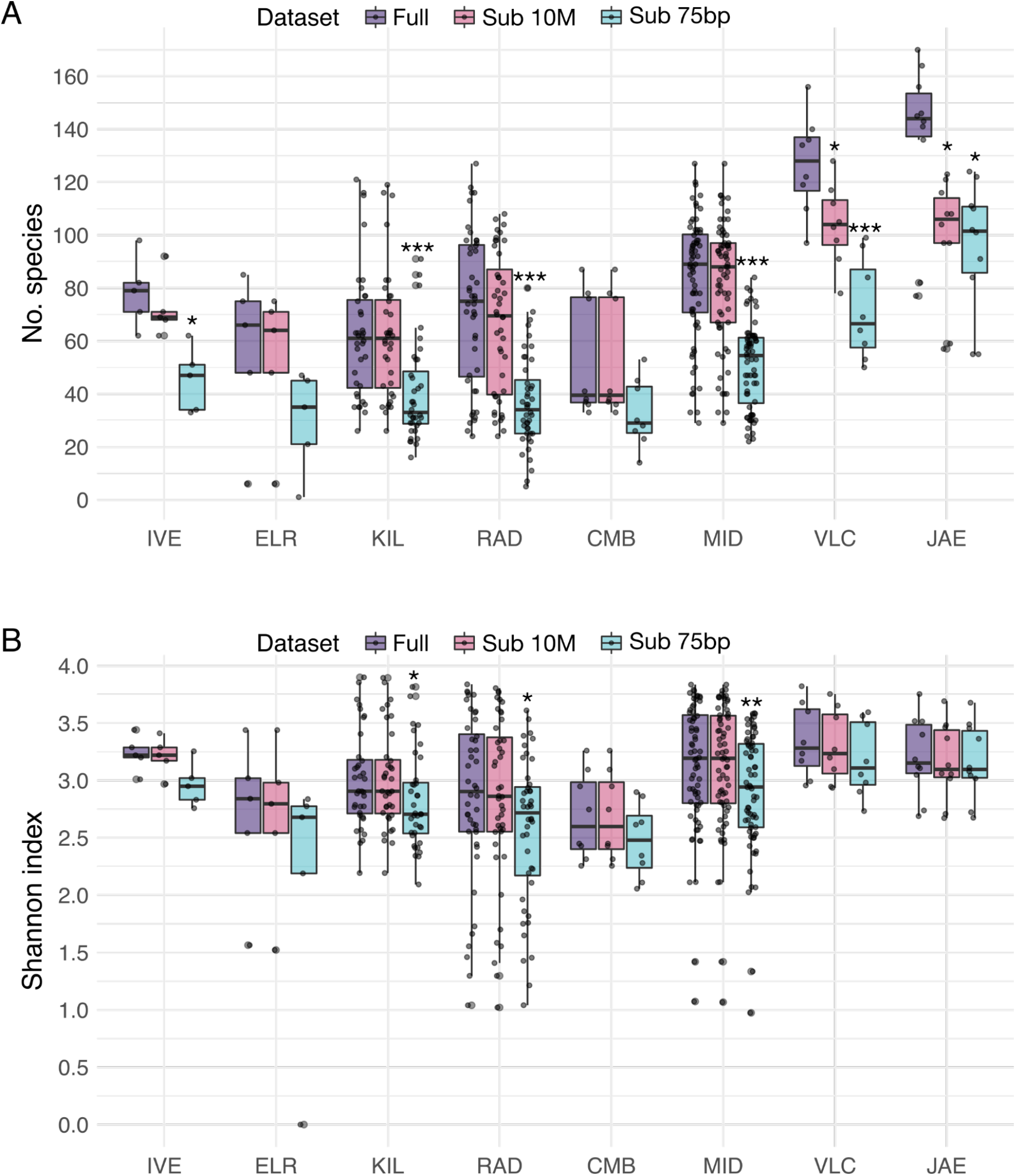
Diversity indices for full and sub-sampled libraries. **A.** Number of species. **B.** Shannon index. *** p < 0.001, ** p < 0.01, * p < 0.05 compared to the full dataset. **Sub 10M** - libraries subsampled to include no more than 10 million reads. **Sub 75bp** - libraries subsampled to include only reads ≤ 75bp in length

**Figure S11.**
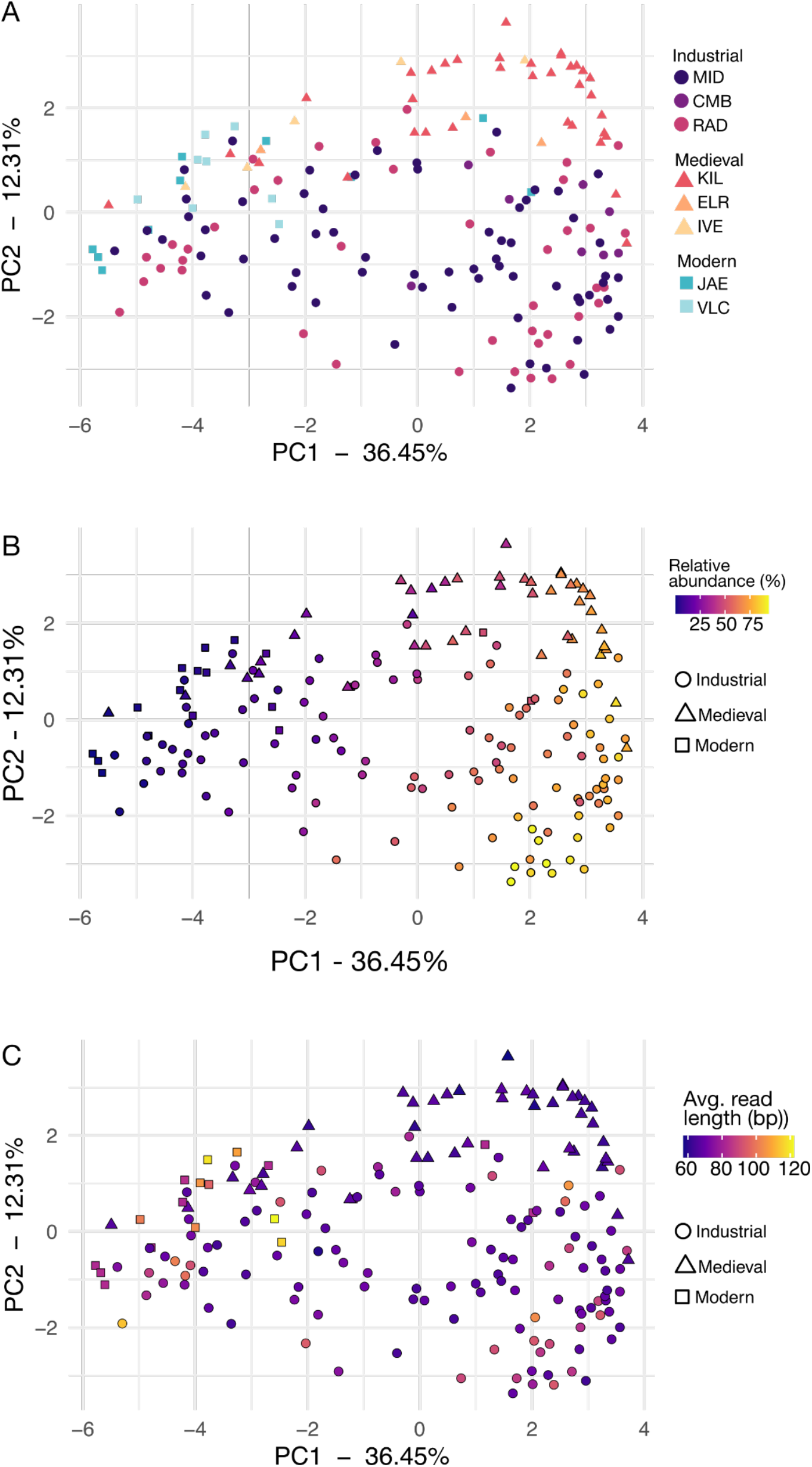
Beta-diversity PCA plot of calculus samples from pre- and post-introduction of tobacco to Europe. Shapes in both indicate time period. These are the same plot as main figure XX5 but colored by **A.** Site. **B.** Relative abundance of the 10 species with strongest positive PC1 loadings, all of which are anaerobic species found late in dental biofilm development. See Supplemental table SXX4. **C.** Average read length per sample.

**Table S4.** Top 10 species with strongest loadings in PC1 in the PCA with pre- and post-tobacco introduction to Europe samples (Figure XX5, Supplemental Figure XX8).

| PC1 direction | Species* | Aerotolerance |
| --- | --- | --- |
| Positive | *Eubacterium minutum* | Anaerobic |
| Positive | Anaerolineaceae bacterium oral taxon 439 | Anaerobic |
| Positive | *Methanobrevibacter oralis* | Anaerobic |
| Positive | *Desulfobulbus oralis* | Anaerobic |
| Positive | *Fretibacterium fastidiosum* | Anaerobic |
| Positive | *Desulfomicrobium orale* | Anaerobic |
| Positive | *Eubacterium saphenum* | Anaerobic |
| Positive | *Peptostreptococcaceae* bacterium oral taxon 113 | Unclear |
| Positive | *Tannerella forsythia* | Anaerobic |
| Positive | *Treponema socranskii* | Anaerobic |
| Negative | *Streptococcus sanguinis* | Facultative |
| Negative | *Lautropia mirabilis* | Facultative |
| Negative | *Neisseria sicca* | Aerobic |
| Negative | *Neisseria mucosa* | Aerobic |
| Negative | *Ottowia* sp oral taxon 894 | Unclear |
| Negative | *Neisseria elongata* | Aerobic |
| Negative | *Capnocytophaga sputigena* | Facultative |
| Negative | *Capnocytophaga gingivalis* | Facultative |
| Negative | *Rothia aeria* | Aerobic |
| Negative | *Streptococcus oralis* | Facultative |

\* Species are ordered from strongest to weakest loading.

**Figure S12.**
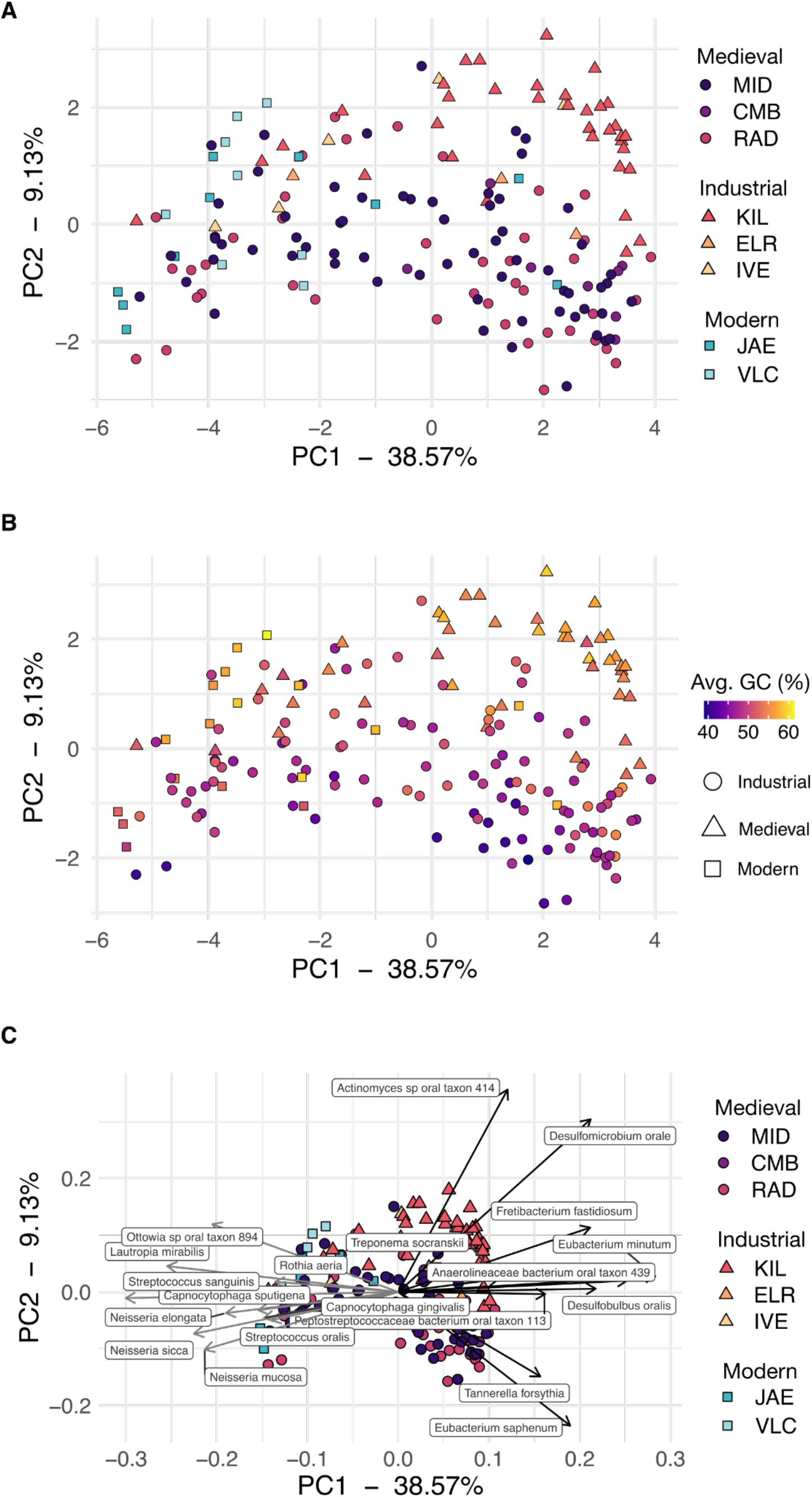
PCA of the species table excluding *Methanobrevibacter oralis*. Removal of this species only slightly alters the relationship of samples in the PCA. **A.** PCA with points colored by site and shaped by time period. **B.** Same as A with points colored by sample average GC content. **C.** Same as A with a biplot showing the top 10 species with strongest PC1 positive and negative loadings. Site codes: **MID** - Middenbeemster, **CMB** - Convento de los Mercedarios de Burtzeña, **RAD** - Radcliffe, **KIL** - Kilteasheen, **ELR** - El Raval, **IVE** - Iglesia de la Virgen de la Estrella, **JAE** - Jaen, **VLC** - Valencia.

**Figure S13.**
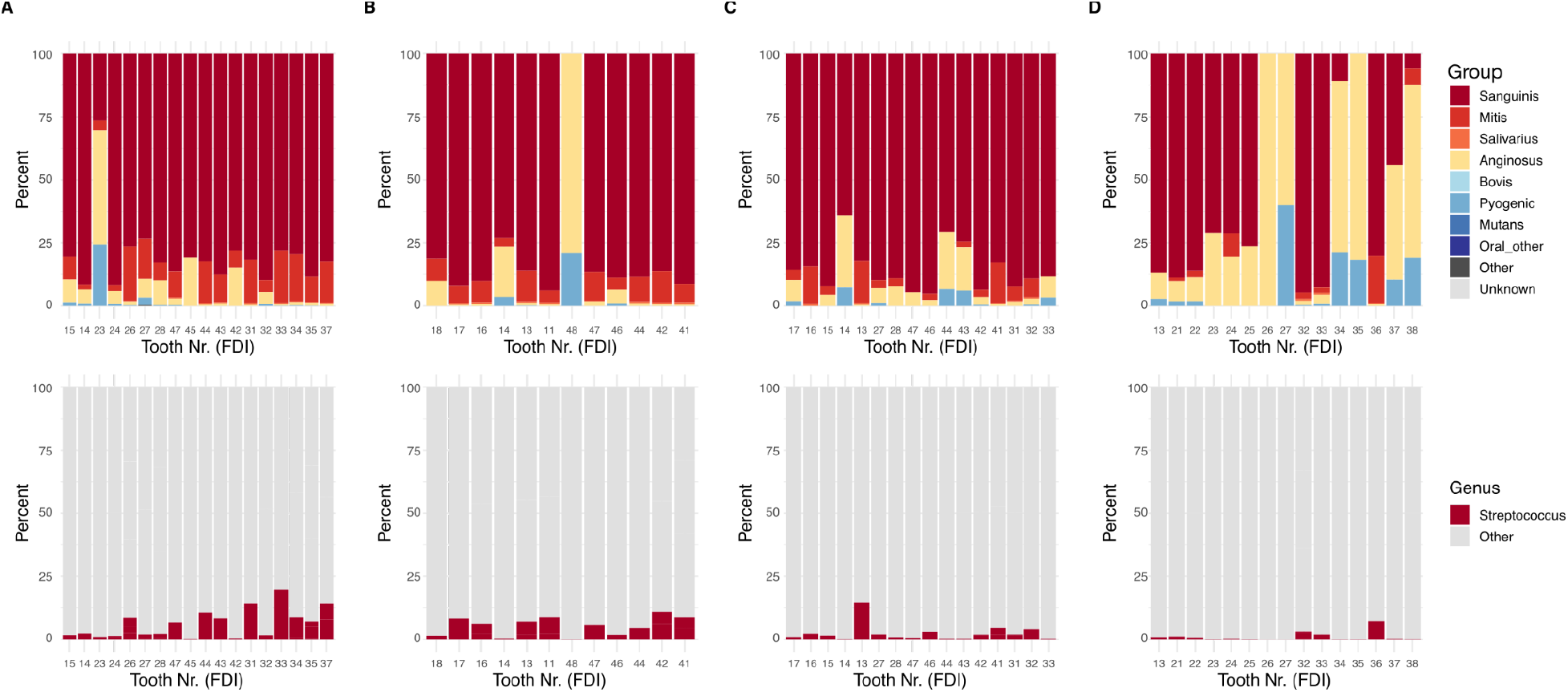
Distribution of *Streptococcus* groups (top panels) and proportion of *Streptococcus* reads (bottom panels) in individual teeth of 4 individuals from Camino del Molino, Spain. **A.** Individual CM55, **B.** Individual CM59, **C.** Individual CM82, **D.** Individual CM165. In cases where multiple surfaces of a tooth were sampled, the proportions were averaged across all surfaces to create a tooth-wide average, which was plotted. Tooth Nr. (FDI) - tooth number based on the FDI World Dental Federation notation.

## References

Abusleme L, Hoare A, Hong B-Y, Diaz PI. 2021. Microbial signatures of health, gingivitis, and periodontitis. Periodontol 2000 86:57–78.

Adler CJ, Dobney K, Weyrich LS, Kaidonis J, Walker AW, Haak W, Bradshaw CJA, Townsend G, Sołtysiak A, Alt KW, Parkhill J, Cooper A. 2013. Sequencing ancient calcified dental plaque shows changes in oral microbiota with dietary shifts of the Neolithic and Industrial revolutions. Nat Genet 45:450–5, 455e1.

Ahlmann-Eltze C, Patil I. 2021. ggsignif: R Package for Displaying Significance Brackets for “ggplot2.” PsyArXiv. doi:10.31234/osf.io/7awm6

Albandar JM, Streckfus CF, Adesanya MR, Winn DM. 2000. Cigar, Pipe, and Cigarette Smoking as Risk Factors for Periodontal Disease and Tooth Loss. Journal of Periodontology. doi:10.1902/jop.2000.71.12.1874

Al Bataineh MT, Dash NR, Elkhazendar M, Alnusairat DMH, Darwish IMI, Al-Hajjaj MS, Hamid Q. 2020. Revealing oral microbiota composition and functionality associated with heavy cigarette smoking. J Transl Med 18:421.

Al-Zahrani MS, Alhassani AA, Melis M, Zawawi KH. 2021. Depression is related to edentulism and lack of functional dentition: An analysis of NHANES data, 2005-2016. J Public Health Dent 81:206–213.

Andrews S. 2010. FastQC: a quality control tool for high throughput sequence data.

Aron F, Brandt G. 2020. Amplification and Pooling v1. protocols.io. doi:10.17504/protocols.io.beqkjduw

Aron F, Hofman C, Fagernäs Z, Velsko I, Skourtanioti E, Brandt G, Warinner C. 2020a. Ancient DNA extraction from dental calculus v1. protocols.io. doi:10.17504/protocols.io.bidyka7w

Aron F, U Neumann G, Brandt G. 2020b. Half-UDG treated double-stranded ancient DNA library preparation for Illumina sequencing v1. protocols.io. doi:10.17504/protocols.io.bmh6k39e

Aten D, Bossaers K, Dehé J, Kuypershoek E, Misset C, Schaap E, Steenhuis M, Van der Wiel K. 2012. 400 jaar Beemster 1612-2012. Wormerveer: Stichting Uitgeverij Noord-Holland.

Axelsson P, Paulander J, Lindhe J. 1998. Dental status and smoking habits among 35-75 year olds in Sweden. Journal of Clinical Periodontal 25:297–305.

Baetsen S, Weterings-Korthorst L. 2013. De menselijke overblijfselen In: Arts N, editor. Een Knekelveld Maakt Geschiedenis, Het Archeologisch Onderzoek Vanhet Koor En Het Grafveld van de Middeleeuwse Catharinakerk in Eindhoven, circa 1200-1850. Matrijs: Utrecht. pp. 151–212.

Baljoon M, Natto S, Abanmy A, Bergstrom J. 2005. Smoking and vertical bone defects in a Saudi Arabian population. Oral Health Prev Dent 3:173.

Becker RA, Wilks AR, Brownrigg R, Minka TP, Deckmyn A. 2021. Maps: Draw geographical maps. Comprehensive R Archive Network (CRAN). https://CRAN.R-project.org/package=maps

Beghini F, McIver LJ, Blanco-Míguez A, Dubois L, Asnicar F, Maharjan S, Mailyan A, Manghi P, Scholz M, Thomas AM, Valles-Colomer M, Weingart G, Zhang Y, Zolfo M, Huttenhower C, Franzosa EA, Segata N. 2021. Integrating taxonomic, functional, and strain-level profiling of diverse microbial communities with bioBakery 3. Elife 10:e65088.

Beghini F, Renson A, Zolnik CP, Geistlinger L, Usyk M, Moody TU, Thorpe L, Dowd JB, Burk R, Segata N, Jones HE, Waldron L. 2019. Tobacco exposure associated with oral microbiota oxygen utilization in the New York City Health and Nutrition Examination Study. Ann Epidemiol 34:18–25.e3.

Bergstrom J. 2014. Smoking rate and periodontal disease prevalence: 40-year trends in Sweden 1970-2010. J Clin Periodontol 41:952–957.

Bergström J. 2005. Tobacco smoking and subgingival dental calculus. J Clin Periodontol 32:81–88.

Bivand R, Rundel C, Pebesma E. 2017. rgeos: interface to geometry engine-open source (GEOS). R package version 0 3–26.

Brealey JC, Leitão HG, van der Valk T, Xu W, Bougiouri K, Dalén L, Guschanski K. 2020. Dental Calculus as a Tool to Study the Evolution of the Mammalian Oral Microbiome. Mol Biol Evol 37:3003–3022.

Briscoe L, Balliu B, Sankararaman S, Halperin E, Garud NR. 2022. Evaluating supervised and unsupervised background noise correction in human gut microbiome data. PLoS Comput Biol 18:e1009838.

Brongers GA. 1964. Nicotiana tabacum: the history of tobacco and tobacco smoking in the Netherlands. Becht.

Brooks S, Suchey JM. 1990. Skeletal age determination based on the os pubis: A comparison of the Acsádi-Nemeskéri and Suchey-Brooks methods. Hum Evol 5:227–238.

Brothwell DR. 1981. Digging Up Bones: The Excavation, Treatment, and Study of Human Skeletal Remains. Cornell University Press.

Buckberry JL, Chamberlain AT. 2002. Age estimation from the auricular surface of the ilium: a revised method. Am J Phys Anthropol 119:231–239.

Buduneli N. 2021. Environmental factors and periodontal microbiome. Periodontol 2000 85:112–125.

Buikstra JE, Ubelaker DH. 1995. Standards for Data Collection from Human Skeletal Remains. The Quarterly Review of Biology. doi:10.1086/419244

Clevis H, Constandse-Westermann T. 1992. De doden vertellen. Opgraving in de Broerenkerk te Zwolle 1987-88. Stichting Archaeologie Ijssel/Vechtstreetk: Kampen.

Cortes Bárcena C. 2013. Dutch and English clay pipes found in Santander (Cantabria, Spain).Journal of the Académie Internationale de la Pipe 6:83–85.

Danecek P, Bonfield JK, Liddle J, Marshall J, Ohan V, Pollard MO, Whitwham A, Keane T, McCarthy SA, Davies RM, Li H. 2021. Twelve years of SAMtools and BCFtools. Gigascience 10. doi:10.1093/gigascience/giab008

Davis NM, Proctor DM, Holmes SP, Relman DA, Callahan BJ. 2018. Simple statistical identification and removal of contaminant sequences in marker-gene and metagenomics data. Microbiome 6:226.

de Heredia Bercero JB, Alaix NM i., Soberón M. 2012. Production and Trade of short stemmed Clay Pipes Found in Barcelona between the Seventeenth and the Nineteenth Century. Journal of the Academie Internationale De La Pipe 5.

de Jong R, Mit ME, Pielage GJ, Haartsen AJ. 1988. Nominatiedossier (nederlandse versie): droogmakerij de Beemster aan de hand waarvan de UNESCO droogmakerij de Beemster op 1 december 1999 op de werelderfgoedlijst heeft geplaatst. Netherlands Department for Conservation, Zeist.

de Miguel Ibáñez MP. 2007. Anexo. Necrópolis mudéjar de Crevillent: estudio osteoarqueológico. Lucentum 221–231.

Domínguez Ballesteros E, Zufiaurre LS, Collado MIG. 2018. El monasterio mercedario de Burtzeña. Kobie Paleoantropología 185–198.

Duran-Pinedo AE, Chen T, Teles R, Starr JR, Wang X, Krishnan K, Frias-Lopez J. 2014. Community-wide transcriptome of the oral microbiome in subjects with and without periodontitis. ISME J 8:1659–1672.

Eke PI, Dye BA, Wei L, Slade GD, Thornton-Evans GO, Borgnakke WS, Taylor GW, Page RC, Beck JD, Genco RJ. 2015. Update on Prevalence of Periodontitis in Adults in the United States: NHANES 2009 to 2012. J Periodontol 86:611–622.

Ewels P, Magnusson M, Lundin S, Käller M. 2016. MultiQC: summarize analysis results for multiple tools and samples in a single report. Bioinformatics 32:3047–3048.

Fagernäs Z, García-Collado MI, Hendy J, Hofman CA, Speller C, Velsko I, Warinner C. 2020. A unified protocol for simultaneous extraction of DNA and proteins from archaeological dental calculus. J Archaeol Sci 118:105135.

Fagernäs Z, Salazar-García DC, Fernández AA, Uriarte MH, Henry A, Maurandi JL, Ozga A, Velsko IM, Warinner C. 2022. Understanding the microbial biogeography of ancient human dentitions to guide study design and interpretation. FEMS Microbes. doi:10.1093/femsmc/xtac006

Falger VSE, Beemsterboer-Köhne CA, Kölker AJ. 2012. Nieuwe Kroniek van de Beemster. Serendipity.

Fellows Yates JA, Lamnidis TC, Borry M, Valtueña AA, Fagernäs Z, Clayton S, Garcia MU, Neukamm J, Peltzer A. 2021a. Reproducible, portable, and efficient ancient genome reconstruction with nf-core/eager. PeerJ. doi:10.1101/2020.06.11.145615

Fellows Yates JA, Velsko IM, Aron F, Posth C, Hofman CA, Austin RM, Parker CE, Mann AE, Nägele K, Arthur KW, Arthur JW, Bauer CC, Crevecoeur I, Cupillard C, Curtis MC, Dalén L, Díaz-Zorita Bonilla M, Díez Fernández-Lomana JC, Drucker DG, Escribano Escrivá E, Francken M, Gibbon VE, González Morales MR, Grande Mateu A, Harvati K, Henry AG, Humphrey L, Menéndez M, Mihailović D, Peresani M, Rodríguez Moroder S, Roksandic M, Rougier H, Sázelová S, Stock JT, Straus LG, Svoboda J, Teßmann B, Walker MJ, Power RC, Lewis CM, Sankaranarayanan K, Guschanski K, Wrangham RW, Dewhirst FE, Salazar-García DC, Krause J, Herbig A, Warinner C. 2021b. The evolution and changing ecology of the African hominid oral microbiome. Proc Natl Acad Sci U S A 118. doi:10.1073/pnas.2021655118

García-Collado MI, Ballesteros ED, Zufiaurre LS. 2018. Estudio osteoarqueológico de la población humana enterrada en el Convento Mercedario de Burtzeña (Barakaldo, Bizkaia), finales s. XVI--principios s. XIX. Kobie Paleoantropología 36:199–222.

Gately I. 2001. La Diva Nicotina: The Story of how Tobacco Seduced the World. Scribner.

Geber J, Murphy E. 2018. Dental markers of poverty: Biocultural deliberations on oral health of the poor in mid-nineteenth-century Ireland. Am J Phys Anthropol 167:840–855.

Goodman J. 1993. Tobacco in history: The cultures of dependence. Routledge.

Granehäll L, Huang KD, Tett A, Manghi P, Paladin A, O’Sullivan N, Rota-Stabelli O, Segata N, Zink A, Maixner F. 2021. Metagenomic analysis of ancient dental calculus reveals unexplored diversity of oral archaeal Methanobrevibacter. Microbiome 9:197.

Hakvoort A. 2013. De begravingen bij de Keyserkerk te Middenbeemster. HOLLANDIA archeologen.

Hardy K, Radini A, Buckley S, Sarig R, Copeland L, Gopher A, Barkai R. 2016. Dental calculus reveals potential respiratory irritants and ingestion of essential plant-based nutrients at Lower Palaeolithic Qesem Cave Israel. Quat Int 398:129–135.

Hendy J, Warinner C, Bouwman A, Collins MJ, Fiddyment S, Fischer R, Hagan R, Hofman CA, Holst M, Chaves E, Klaus L, Larson G, Mackie M, McGrath K, Mundorff AZ, Radini A, Rao H, Trachsel C, Velsko IM, Speller CF. 2018. Proteomic evidence of dietary sources in ancient dental calculus. Proc Biol Sci 285:20180977.

Heng CK, Badner VM, Freeman KD. 2006. Relationship of Cigarette Smoking to Dental Caries in a Population of Female Inmates. J Correct Health Care 12:164–174.

Herbig A, Maixner F, Bos KI, Zink A, Krause J, Huson DH. 2016. MALT: Fast alignment and analysis of metagenomic DNA sequence data applied to the Tyrolean Iceman. bioRxiv. doi:10.1101/050559

Hillson S. 2001. Recording dental caries in archaeological human remains. Int J Osteoarchaeol 11:249–289.

Hoffman GE, Roussos P. 2021. Dream: powerful differential expression analysis for repeated measures designs. Bioinformatics 37:192–201.

Hoffman GE, Schadt EE. 2016. variancePartition: interpreting drivers of variation in complex gene expression studies. BMC Bioinformatics 17:483.

Huang C, Shi G. 2019. Smoking and microbiome in oral, airway, gut and some systemic diseases. J Transl Med 17:225.

Hughes J. 2003. Learning to Smoke: Tobacco Use in the West. University of Chicago Press.

Huson DH, Beier S, Flade I, Górska A, El-Hadidi M, Mitra S, Ruscheweyh H-J, Tappu R. 2016. MEGAN Community Edition - Interactive Exploration and Analysis of Large-Scale Microbiome Sequencing Data. PLoS Comput Biol 12:e1004957.

Inskip SA, Zachary L, Serrano Ruber M, Hoogland MLP. n.d. Pipe Smoking and Oral Health in Males from the Netherlands during the 18th-19th century. Post Medieval Archaeology.

Işcan MY, Loth SR. 1986. Determination of age from the sternal rib in white females: a test of the phase method. J Forensic Sci 31:990–999.

Jackes M. 2015. The Zwolle teeth: an independent look at the data.

Jersie-Christensen RR, Lanigan LT, Lyon D, Mackie M, Belstrøm D, Kelstrup CD, Fotakis AK, Willerslev E, Lynnerup N, Jensen LJ, Cappellini E, Olsen JV. 2018. Quantitative metaproteomics of medieval dental calculus reveals individual oral health status. Nat Commun 9:4744.

Kassambara A. 2020. rstatix: pipe-friendly framework for basic statistical tests. R package version 0.4. 0.

Kazarina A, Petersone-Gordina E, Kimsis J, Kuzmicka J, Zayakin P, Griškjans Ž, Gerhards G, Ranka R. 2021. The Postmedieval Latvian Oral Microbiome in the Context of Modern Dental Calculus and Modern Dental Plaque Microbial Profiles. Genes 12. doi:10.3390/genes12020309

Knights D, Kuczynski J, Charlson ES, Zaneveld J, Mozer MC, Collman RG, Bushman FD, Knight R, Kelley ST. 2011. Bayesian community-wide culture-independent microbial source tracking. Nat Methods 8:761–763.

Kolenbrander PE, Palmer RJ Jr, Rickard AH, Jakubovics NS, Chalmers NI, Diaz PI. 2006. Bacterial interactions and successions during plaque development. Periodontol 2000 42:47–79.

Kowalski CJ. 1971. Relationship between smoking and calculus deposition. J Dent Res 50:101–104.

Kumar PS, Matthews CR, Joshi V, de Jager M, Aspiras M. 2011. Tobacco smoking affects bacterial acquisition and colonization in oral biofilms. Infect Immun 79:4730–4738.

Kvaal SI, Derry TK. 1996. Tell-tale teeth: abrasion from the traditional clay pipe. Endeavour 20:28–30.

Lee H, Kim D, Jung A, Chae W. 2022. Ethnicity, social, and clinical risk factors to tooth loss among older adults in the U.s., NHANES 2011-2018. Int J Environ Res Public Health 19:2382.

Leroux H. 2012. The use of dental nonmetric traits for intracemetery kinship analysis and cemetery structure analysis from the site of Middenbeemster, the Netherlands. University of Leiden.

Listgarten MA, Mayo H, Amsterdam M. 1973. Ultrastructure of the attachment device between coccal and filamentous microorganisms in “corn cob” formations of dental plaque. Arch Oral Biol 18:651–IN5.

Listgarten MA, Mayo HE, Tremblay R. 1975. Development of dental plaque on epoxy resin crowns in man. A light and electron microscopic study. J Periodontol 46:10–26.

Lovejoy CO, Meindl RS, Pryzbeck TR, Mensforth RP. 1985. Chronological metamorphosis of the auricular surface of the ilium: a new method for the determination of adult skeletal age at death. Am J Phys Anthropol 68:15–28.

Lukacs JR, Largaespada LL. 2006. Explaining sex differences in dental caries prevalence: saliva, hormones, and “life-history” etiologies. Am J Hum Biol 18:540–555.

Lukacs Jr. 2008. Fertility and Agriculture Accentuate Sex Differences in Dental Caries Rates. Curr Anthropol 49:901–914.

Maat GJ. 2001. Diet and age-at-death determinations from molar attrition. A review related to the Low Countries J Forensic Odonto-Stomatol 19:18–21.

Mann AE, Sabin S, Ziesemer K, Vågene ÅJ, Schroeder H, Ozga AT, Sankaranarayanan K, Hofman CA, Fellows Yates JA, Salazar-García DC, Frohlich B, Aldenderfer M, Hoogland M, Read C, Milner GR, Stone AC, Lewis CM Jr, Krause J, Hofman C, Bos KI, Warinner C. 2018. Differential preservation of endogenous human and microbial DNA in dental calculus and dentin. Sci Rep 8:9822.

Martí JT, Pérez JRO, Gómez IR, Bebia MAE. 2009. El cementerio mudéjar del Raval (Crevillent-Alicante). AyTM 16:179–216.

Martinez-Canut P, Lorca A, Magán R. 1995. Smoking and periodontal disease severity. J Clin Periodontol 22:743–749.

Mason MR, Preshaw PM, Nagaraja HN, Dabdoub SM, Rahman A, Kumar PS. 2015. The subgingival microbiome of clinically healthy current and never smokers. ISME J 9:268–272.

Moon J-H, Lee J-H, Lee J-Y. 2015. Subgingival microbiome in smokers and non-smokers in Korean chronic periodontitis patients. Mol Oral Microbiol 30:227–241.

Morton JT, Toran L, Edlund A, Metcalf JL, Lauber C, Knight R. 2017. Uncovering the Horseshoe Effect in Microbial Analyses. mSystems 2. doi:10.1128/mSystems.00166-16

Nociti FH Jr, Casati MZ, Duarte PM. 2015. Current perspective of the impact of smoking on the progression and treatment of periodontitis. Periodontol 2000 67:187–210.

Norton M. 2008. Sacred Gifts, Profane, Pleasures: A History of Tobacco and Chocolate in the Atlantic World. Cornell University Press.

Ogden A. 2007. Advances in the palaeopathology of teeth and jawsAdvances in Human Palaeopathology. Chichester, UK: John Wiley & Sons, Ltd. pp. 283–307.

Oksanen, J. 2007. Vegan : community ecology package version 1.8-6. http://cran.r-project.org.

Ottoni C, Borić D, Cheronet O, Sparacello V, Dori I, Coppa A, Antonović D, Vujević D, Price TD, Pinhasi R, Cristiani E. 2021. Tracking the transition to agriculture in Southern Europe through ancient DNA analysis of dental calculus. Proc Natl Acad Sci U S A 118. doi:10.1073/pnas.2102116118

Passalacqua NV. 2009. Forensic Age-at-Death Estimation from the Human Sacrum. Journal of Forensic Sciences. doi:10.1111/j.1556-4029.2008.00977.x

Pebesma E. 2018. Simple features for R: Standardized support for spatial vector data. R J 10:439.

Pedersen TL. 2017. patchwork: The Composer of ggplots. R package version 0 0 1.

Pejčić A, Obradović R, Kesić L, Kojović D. 2007. Smoking and periodontal disease: A review. Med Biol 14:53–59.

Phenice TW. 1969. A newly developed visual method of sexing the os pubis. Am J Phys Anthropol 30:297–301.

Radini A, Tromp M, Beach A, Tong E, Speller C, McCormick M, Dudgeon JV, Collins MJ, Rühli F, Kröger R, Warinner C. 2019. Medieval women’s early involvement in manuscript production suggested by lapis lazuli identification in dental calculus. Sci Adv 5:eaau7126.

Richards VP, Palmer SR, Pavinski Bitar PD, Qin X, Weinstock GM, Highlander SK, Town CD, Burne RA, Stanhope MJ. 2014. Phylogenomics and the dynamic genome evolution of the genus Streptococcus. Genome Biol Evol 6:741–753.

Rohart F, Gautier B, Singh A, Lê Cao K-A. 2017. mixOmics: An R package for ‘omics feature selection and multiple data integration. PLoS Comput Biol 13:e1005752.

Salazar-García DC, Power RC, Rudaya N, Kolobova K, Markin S, Krivoshapkin A, Henry AG, Richards MP, Viola B. 2021. Dietary evidence from Central Asian Neanderthals: A combined isotope and plant microremains approach at Chagyrskaya Cave (Altai, Russia). J Hum Evol 156:102985.

Salazar-García DC, Richards MP, Nehlich O, Henry AG. 2014. Dental calculus is not equivalent to bone collagen for isotope analysis: a comparison between carbon and nitrogen stable isotope analysis of bulk dental calculus, bone and dentine collagen from same individuals from the Medieval site of El Raval (Alicante, Spain). Journal of Archaeological Science. doi:10.1016/j.jas.2014.03.026

Schabbink ML. 2020. Een Beemster Poldermolen. Archaeologisch onderzoek rond de Draaioorder molengang in Zuidoostbeemster (No. 6). Noord-Hollandse Archaeologisch Publicaties - 9. Huis Van Hilde Archaeologiecentrum Noord-Holland: Castricum.

Schmid C. 2022. ggpointgrid: Rearrange scatter plot points on a regular grid. https://github.com/nevrome/ggpointgrid

Socransky SS, Manganiello AD, Propas D, Oram V, van Houte J. 1977. Bacteriological studies of developing supragingival dental plaque. J Periodontal Res 12:90–106.

South A. 2017a. rnaturalearth: World map data from Natural Earth. R package version 0.1. 0. The R Foundation https://CRANR-project.org/package=rnaturalearth.

South A. 2017b. rnaturalearthdata: world vector map data from Natural Earth used in’rnaturalearth’. R package version 0.1. 0.

Sreedevi M, Ramesh A, Dwarakanath C. 2012. Periodontal status in smokers and nonsmokers: a clinical, microbiological, and histopathological study. Int J Dent 2012:571590.

Stahl R, Warinner C, Velsko I, Orfanou E, Aron F, Brandt G. 2021. Illumina double-stranded DNA dual indexing for ancient DNA v2. protocols.io. doi:10.17504/protocols.io.bvt8n6rw

Stam RD. 2019. Vergeten glorie: De economische ontwikkeling van de Nederlandse kleipijpennijverheid in de 17e en 18e eeuw, met speciale aandacht voor de export. Universiteit Utrecht.

Tamashiro R, Strange L, Schnackenberg K, Santos J, Gadalla H, Zhao L, Li EC, Hill E, Hill B, Sidhu G, Kirst M, Walker C, Wang GP. 2021. Stability of healthy subgingival microbiome across space and time. Sci Rep 11:23987.

Theilade E, Fejerskov O, Prachyabrued W, Kilian M. 1974. Microbiologic study on developing plaque in human fissures. Scand J Dent Res 82:420–427.

Theilade E, Wright WH, Jensen SB, Löe H. 1966. Experimental gingivitis in man. II. A longitudinal clinical and bacteriological investigation. J Periodontal Res 1:1–13.

Tinanoff N, Gross A, Brady JM. 1976. Development of plaque on enamel. Parallel investigations. J Periodontal Res 11:197–209.

Trelis J, Ortega JR, Reina I, Esquembre MA. 2010. El Cementeri Mudèjar del Raval (Crevillent-Alacant). Rella, La 213–252.

Tullett W. 2019. Smell in Eighteenth-Century England: A Social Sense. Oxford University Press.

Vågene ÅJ, Herbig A, Campana MG, Robles García NM, Warinner C, Sabin S, Spyrou MA, Andrades Valtueña A, Huson D, Tuross N, Bos KI, Krause J. 2018. Salmonella enterica genomes from victims of a major sixteenth-century epidemic in Mexico. Nat Ecol Evol 2:520–528.

Veldman JK. 2013. Non-metric traits. An assessment of cranial and post-cranial non-metric traits in the skeletal assemblage from the 17th-19th century churchyard of middenbeemster, the Netherlands. University of Leiden.

Vellappally S, Fiala Z, Smejkalová J, Jacob V, Shriharsha P. 2007. Influence of tobacco use in dental caries development. Cent Eur J Public Health 15:116–121.

Velsko IM, Fellows Yates JA, Aron F, Hagan RW, Frantz LAF, Loe L, Martinez JBR, Chaves E, Gosden C, Larson G, Warinner C. 2019. Microbial differences between dental plaque and historic dental calculus are related to oral biofilm maturation stage. Microbiome 7:102.

Velsko IM, Frantz LAF, Herbig A, Larson G, Warinner C. 2018. Selection of Appropriate Metagenome Taxonomic Classifiers for Ancient Microbiome Research. mSystems 3. doi:10.1128/mSystems.00080-18

Velsko IM, Overmyer KA, Speller C, Klaus L, Collins MJ, Loe L, Frantz LAF, Sankaranarayanan K, Lewis CM Jr, Martinez JBR, Chaves E, Coon JJ, Larson G, Warinner C. 2017. The dental calculus metabolome in modern and historic samples. Metabolomics 13:134.

Veselka B. 2016. Begraven bij de Sint-Janskerk: analyse van de zeventiende tot negentiende-eeuwse skeletten uit RoosendaalJaarboek de Gulden Roos. pp. 33–44.

Wade WG. 2021. Resilience of the oral microbiome. Periodontol 2000. doi:10.1111/prd.12365

Walker D, Henderson M. 2010. Smoking and health in London’s East End in the first half of the 19th century. Post-Medieval Archaeology 44:209–222.

Warinner C, Rodrigues JFM, Vyas R, Trachsel C, Shved N, Grossmann J, Radini A, Hancock Y, Tito RY, Fiddyment S, Speller C, Hendy J, Charlton S, Luder HU, Salazar-García DC, Eppler E, Seiler R, Hansen LH, Castruita JAS, Barkow-Oesterreicher S, Teoh KY, Kelstrup CD, Olsen JV, Nanni P, Kawai T, Willerslev E, von Mering C, Lewis CM Jr, Collins MJ, Gilbert MTP, Rühli F, Cappellini E. 2014. Pathogens and host immunity in the ancient human oral cavity. Nat Genet 46:336–344.

Wei T, Simko V, Levy M, Xie Y, Jin Y, Zemla J. 2013. corrplot: Visualization of a correlation matrix. R package version 0 73 230.

Western G, Bekvalac J. 2020. Manufactured Bodies: The Impact of Industrialisation on London Health. Oxbow Books.

Wibowo MC, Yang Z, Borry M, Hübner A, Huang KD, Tierney BT, Zimmerman S, Barajas-Olmos F, Contreras-Cubas C, García-Ortiz H, Martínez-Hernández A, Luber JM, Kirstahler P, Blohm T, Smiley FE, Arnold R, Ballal SA, Pamp SJ, Russ J, Maixner F, Rota-Stabelli O, Segata N, Reinhard K, Orozco L, Warinner C, Snow M, LeBlanc S, Kostic AD. 2021. Reconstruction of ancient microbial genomes from the human gut. Nature 594:234–239.

Wilke CO. 2020. cowplot: streamlined plot theme and plot annotations for “ggplot2”. R package version 0.9. 2; 2017. https://cloud.r-project.org/web/packages/cowplot/index.html

Workshop of European Anthropologists. 1980. Recommendations for age and sex diagnoses of skeletons. J Hum Evol 9:517–549.

Yang Y, Zheng W, Cai Q-Y, Shrubsole MJ, Pei Z, Brucker R, Steinwandel MD, Bordenstein SR, Li Z, Blot WJ, Shu X-O, Long J. 2019. Cigarette smoking and oral microbiota in low-income and African-American populations. J Epidemiol Community Health 73:1108–1115.

Yost S, Duran-Pinedo AE, Krishnan K, Frias-Lopez J. 2017. Potassium is a key signal in host-microbiome dysbiosis in periodontitis. PLoS Pathog 13:e1006457.

Yost S, Duran-Pinedo AE, Teles R, Krishnan K, Frias-Lopez J. 2015. Functional signatures of oral dysbiosis during periodontitis progression revealed by microbial metatranscriptome analysis. Genome Med 7:27.

Zee KY, Samaranayake LP, Attström R. 1997. Scanning electron microscopy of microbial colonization of “rapid” and “slow” dental-plaque formers in vivo. Arch Oral Biol 42:735–742.

Zee K-Y, Samaranayake LP, Attstrom R. 1996. Predominant cultivable supragingival plaque in Chinese “rapid” and “slow” plaque formers. Journal of Clinical Periodontology. doi:10.1111/j.1600-051x.1996.tb00532.x

